# Human plasma cells engineered to secrete bispecifics drive effective *in vivo* leukemia killing

**DOI:** 10.1101/2023.08.24.554523

**Authors:** Tyler F. Hill, Parnal Narvekar, Gregory Asher, Nathan Camp, Kerri R. Thomas, Sarah K. Tasian, David J. Rawlings, Richard G. James

## Abstract

Bispecific antibodies are an important tool for the management and treatment of acute leukemias. Advances in genome-engineering have enabled the generation of human plasma cells that secrete therapeutic proteins and are capable of long-term *in vivo* engraftment in humanized mouse models. As a next step towards clinical translation of engineered plasma cells (ePCs) towards cancer therapy, here we describe approaches for the expression and secretion of bispecific antibodies by human plasma cells. We show that human ePCs expressing either fragment crystallizable domain deficient anti-CD19 x anti-CD3 (blinatumomab) or anti-CD33 x anti-CD3 bispecific antibodies mediate T cell activation and direct T cell killing of specific primary human cell subsets and B-acute lymphoblastic leukemia or acute myeloid leukemia cell lines *in vitro*. We demonstrate that knockout of the self-expressed antigen, CD19, boosts anti-CD19 bispecific secretion by ePCs and prevents self-targeting. Further, anti-CD19 bispecific-ePCs elicited tumor eradication *in vivo* following local delivery in flank-implanted Raji lymphoma cells. Finally, immunodeficient mice engrafted with anti-CD19 bispecific-ePCs and autologous T cells potently prevented *in vivo* growth of CD19^+^ acute lymphoblastic leukemia in patient-derived xenografts. Collectively, these findings support further development of ePCs for use as a durable, local delivery system for the treatment of acute leukemias, and potentially other cancers.

**Key points:**

- Using gene editing, we engineered human plasma cells that secrete functional bispecifics to target leukemia cells expressing CD19 or CD33
- Engineered plasma cells secreting bispecifics suppress patient-derived leukemia in immunodeficient mice

## Introduction

Immunotherapies that recruit cytotoxic T cells to kill cancer cells, such as bispecific antibodies, have played a significant role in the improved survival rates for patients with B-cell acute lymphoblastic leukemia (B-ALL)^1–4^. Blinatumomab is an anti-CD19 x anti-CD3 non-immunoglobulin G-like bispecific antibody (non-IgG-like bispecific; also called a bispecific T cell engager, BiTE™) that received FDA approval in 2014 for the treatment of patients with relapsed/refractory B-ALL ^4,5^. Blinatumomab is now used in multiple B-ALL settings, including frontline therapy, as a bridge to transplantation, consolidation therapy, and as a low toxicity alternative to chemotherapy regimens^6^. A limitation of Blinotumamab^7^ and other non-IgG-like bispecifics^8,9^ is that these molecules lack fragment crystallizable domains and have short half-lives, which necessitate continuous high dose intravenous infusions. Moreover, this intensive regimen can be challenging for patients, especially those with limited hospital access^10,11^. A range of methods have been utilized in an attempt to extend biologic half-life of non-IgG-like bispecifics^12^, including conjugation with small molecules, fragment crystallizable domains, or albumin binding motifs. However, it remains unclear whether these fusion molecules will be effective, lack immunogenicity, and/or overcome the need for multiple continuous high-dose infusions.

We, and others, have explored using engineered plasma cells (ePCs) as a long-term biologic drug delivery platform^13–16^. Engineered B cell populations have been investigated in proof-of-concept studies to deliver biologic drugs to treat protein deficiency diseases^13,17^, viral infections^18–22^, and cancer^15,23^. Based on these observations, we predicted that adoptive transfer of bispecific expressing ePC might mitigate challenges related to both bispecific half-life and high dose systemic toxicity. Plasma cells are uniquely suited to deliver biologics over long periods due to their long lifespan^24^ (half-life is estimated to be 11 to 200 years^25^), and high secretory capacity (up to 10,000 IgG molecules per second^26–28)^, Furthermore, *ex-vivo* generated ePCs resemble endogenous plasma cells, and can stably secrete therapeutically relevant levels of immunoglobulin for greater than one year in hIL6-humanized mice^14^. Because PCs^29,30^ and ePCs^14^ preferentially localize to bone marrow and other tissue microenvironments where progenitor B-ALL cells reside^31^, we predicted that ePCs could harmonize with local bispecific delivery to induce potent anti-leukemia activity.

In this study, we describe a homology-directed repair strategy (HDR) based gene editing strategy for the generation of ePC that produce large quantities of anti-CD19 x anti-CD3 or anti-CD33 x anti-CD3 non-IgG-like bispecifics to target B-ALL or acute myeloid leukemia (AML), respectively. Our combined findings demonstrate that ePCs secreting bispecifics can promote T-cell driven killing of primary human cells, human leukemic cell lines *in vitro,* and patient-derived B-ALL xenografts *in vivo*. Based upon our preclinical results, we propose that ePC strategies could be translated to the clinic for evaluation of bispecific delivery to patients with acute leukemias, and other scenarios where half-life is limiting or local delivery could reduce on-target adverse effects.

## Methods

### B cell culturing and PC differentiation

We isolated B cells from healthy human donors’ PBMCs (Fred Hutchinson Cancer Research Center) using the EasySep Human B cell enrichment kit (Stem Cell Technologies). We obtained >95% purity for B cells defined by CD3 negativity and CD19 positivity. Isolated B cells were cultured in Iscove’s modified Dulbecco’s medium (Gibco), supplemented with 2-mercaptoethanol (55μM) and 10% FBS. For Figure 1, cells were cultured for seven days as described in Hung et al^13^. For experiments in Figure 2-6 cells were cultured as described in Cheng et al^32^. Cell concentrations were kept between 5-15×10^5^ live cells per ml. Cells for *in vivo* experiments were purified via CD3 bead depletion column (Miltenyi) prior to injection.

**Figure 1:**
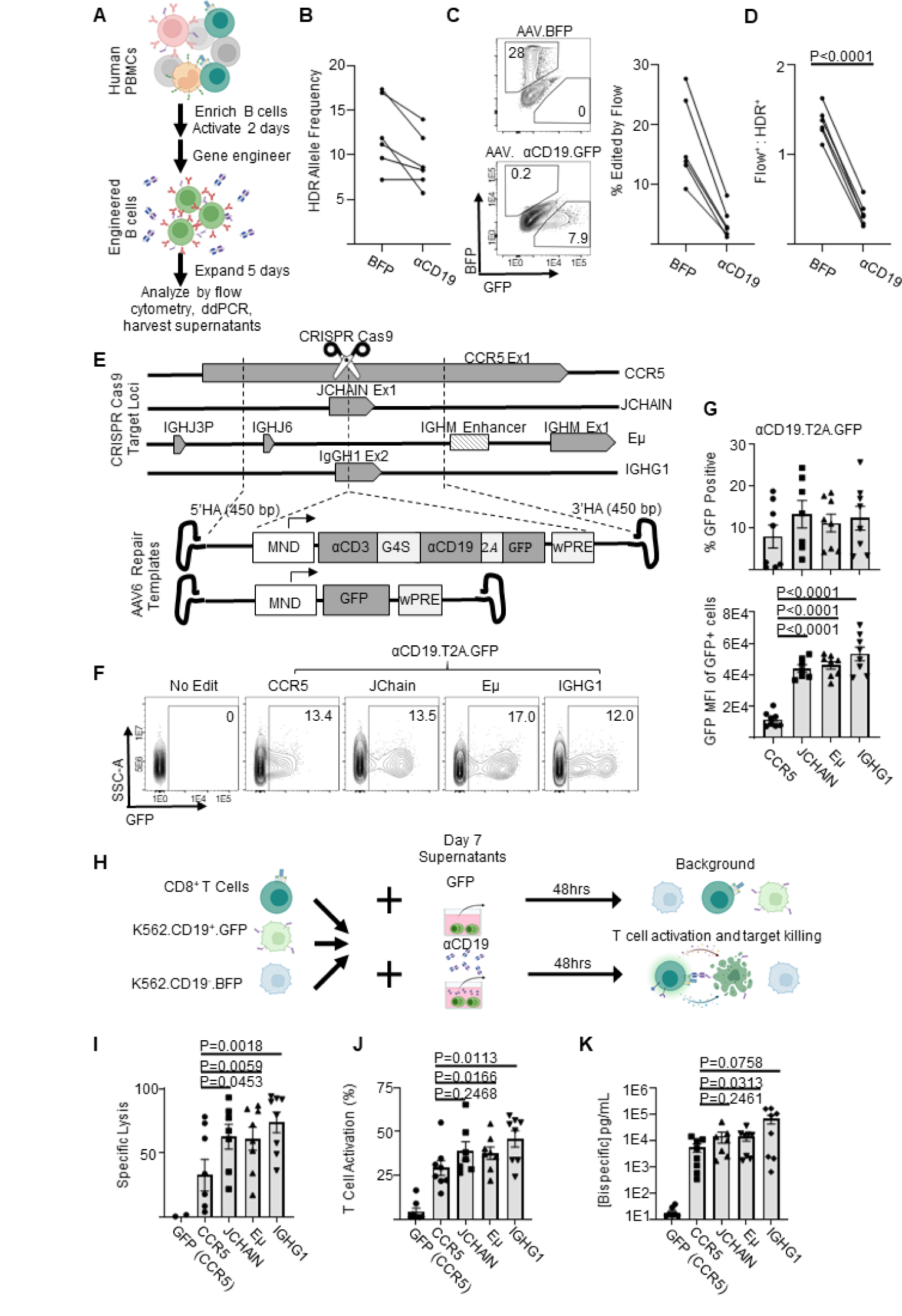
Genome engineered primary human B cells secrete functional αCD19-bispecific in a locus dependent manner. **A)** Schematic showing the experimental flow of a primary B cell experiment. Briefly, after isolation from PBMCs, B cells were edited to express either BFP or αCD19.T2A.GFP transgenes at CCR5 genetic loci via HDR-gene editing with AAV6 delivered DNA repair templates. Five days later genomic DNA, cells and supernatants were analyzed as indicated. **B)** Transgene integration at CCR5 locus shown here as HDR allele frequency was measured by ddPCR. **C)** Representative flow cytometry plots showing transgene expression of fluorescent proteins in engineered B cells shown and quantified as % edited of live cells. **D)** Ratio of engineering rate as determined by ddPCR vs flow cytometry. **E)** Schematic showing the editing strategies for delivery of GFP or αCD19.T2A.GFP to antibody-associated loci. **F)** Representative flow cytometry plots of αCD19.T2A.GFP edited B cells with **G)** the quantification of % edited and GFP mean fluorescent intensity of edited cells. **H)** K562 killing assay schema. Supernatants from edited B cells were incubated with target (CD19^+^) and control (CD19^-^) K562 cells with CD8^+^ T cells for 48 hours. Cells were harvested for flow cytometry to obtain **I)** specific lysis of CD19^+^ K562 and **J)** T cell activation (%CD69^+^CD137^+^ of CD3^+^ cells). **K)** The concentration of bispecific in the supernatants as interpolated from %T cell activated data. Data are from five donors in five independent experiments. Error bars represent SEM. P values calculated using D) a paired student’s t test and G,I-K) paired one-way ANOVAs with Dunnett’s posttest. Illustrations were created in part with biorender.com.

**Figure 2:**
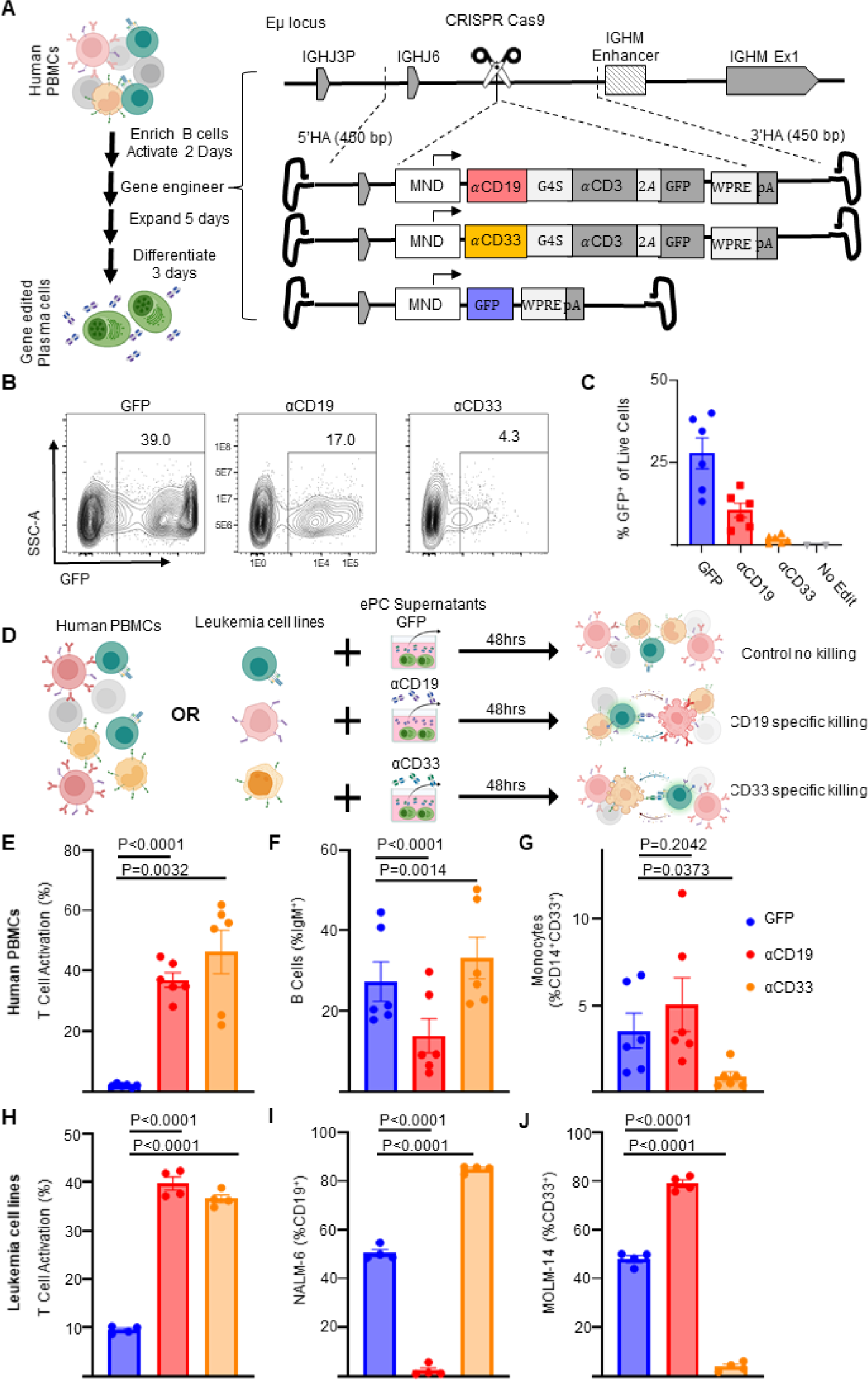
Human plasma cells engineered to secrete anti-leukemia bispecifics specifically target cells expressing physiological levels of antigen. Primary human B cells were isolated and cultured for two days in activating media then edited. **A)** Schematic showing how primary activated human B cells were edited to express GFP or αCD19.T2A.GFP or αCD33.T2A.GFP. After editing activated B cells, the engineered cells were then cultured in expansionary media for 5 days followed by differentiation into PCs over 3 days and cells and supernatants. **B)** Representative flow cytometry plots assessing editing via expression of GFP and **C)** quantification as % of live cells. **D)** Schematic illustrating *in vitro* PBMC or Leukemia cell line killing assays. Briefly, autologous CD8^+^ T cells are co-cultured with PBMCs or mixed leukemia cell populations (NALM-6 and MOLM-14) in the presence of supernatants from ePCs for 48 hours. Flow cytometry was used to quantify **E)** T cell activation (%CD69^+^CD137^+^ of CD3^+^ cells), **F)** the % B cells (IgM^+^) of live cells, **G)** the % monocytes (CD14^+^CD33^+^) of live cells in PBMC cultures at the end of the 48-hour co-culture. Likewise flow cytometry was used to quantify **H)** T cell activation (%CD69^+^,CD137^+^ of CD8^+^ cells), the frequency of **I)** NAML-6 (CD19^+^) and **J)** MOLM-14 (CD33^+^) in the leukemia cell line killing assay. In E-G, data were obtained from six donors in three independent experiments, and in H-J, data were obtained from four donors. Error bars represent SEM. P-values were calculated using paired one-way ANOVAs with Dunnett’s posttest. Illustrations were created in part with biorender.com.

**Figure 3:**
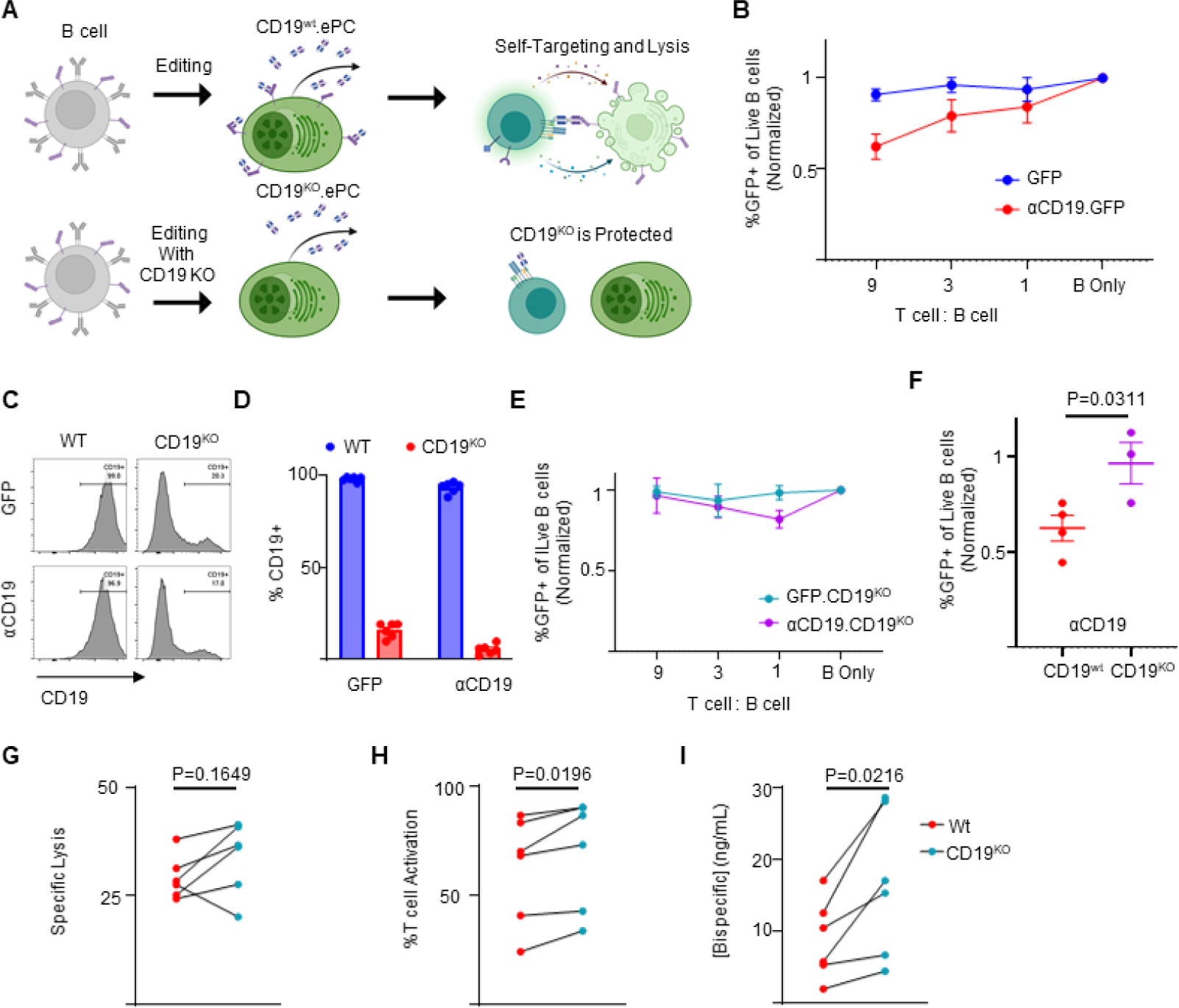
CD19 knockout prevents self-targeting of αCD19-ePCs and increases αCD19-bispecific secretion. **A)** Schematic showing the self-targeting assay of ePCs with and without CD19 knockout. Primary human B cells were engineered to express either GFP or αCD19.T2A.GFP at the Eμ locus, and/or to eliminate CD19. These engineered cells were incubated with the indicated ratios of autologous T cells. **B)** After 24 hours, flow cytometry was used to calculate the percentage of GFP^+^ of live CD20^+^ B cells. The relative quantity of transgene-expressing cells was plotted. sgRNAs targeting CD19 were included to elicit knock out CD19 while engineering into the Eμ. Representative flow cytometry images **C)** and quantification **D)** of CD19 expression in engineered cells is shown. **E)** CD19^KO^ cells were incubated with the indicated ratios of T cells for 24 hours. After incubation of edited cells with T cells, we used flow cytometry to quantify the % GFP^+^ of CD20^+^ cells. **F)** Combined data showing the GFP percentage following incubation of edited cells with T cells at a nine:one ratio. **G-I)** Engineered B cells were further differentiated over 3 days into ePCs. Supernatants from CD19^KO^ and WT αCD19 ePCs were incubated with T cells, K562 CD19^+^ and K562 CD19^-^ cells for 48 hours. **G)** Specific lysis of CD19^+^ K562 and **H)** T cell activation (%CD69^+^CD137^+^ of CD3^+^ cells) was quantified. **I)** αCD19 bispecific concentration was interpolated using recombinant αCD19 bispecific standards curves. These data are from four donors. Error bars represent SEM. P-values were calculated by paired one-way ANOVA with Dunnett’s posttest (F) and paired student’s T test (G-I). Illustrations created in part with biorender.com.

### AAV6 HDR CRISPR Cas9 Engineering of B cells

Clustered regularly interspaced short palindromic repeats (CRISPR) RNAs (crRNAs) targeting the CCR5, JCHAIN, IgG1, Eμ^18^, and CD19 (sequences in Table S1) were identified using the Broad Institute GPP sgRNA Designer (http://portals.broadinstitute.org/gpp/public/analysis-tools/sgrna-design) and synthesized (IDT) containing phosphorothioate linkages and 2′O-methyl modifications. crRNA and trans-activating crRNA (tracrRNA; IDT) single guide hybrids were mixed with 3uM Cas9 nuclease (Berkeley Labs) at a 1.2:1 ratio and delivered to cells by Lonza 3D (CA-137) or Maxcyte GTX (B cell 3) electroporation. After electroporation, cells were transferred into the activation medium (1.5×10^6^cells/mL) in the presence of adeno-associated virus 6 (AAV6) vectors carrying homologous DNA repair templates (20% AAV by volume or viral copy of 1×10^4^ per cell, Figure 1E schema). The medium was changed 24 hours following AAV6 administration. AAV6 vectors were produced as previously described^13^ or manufactured by Sirion Biotech.

### In vitro ePC-mediated killing assays (K562, PBMC, leukemia cell line, and self-killing assays)

For the K562 killing assay, K562 cells were obtained from ATCC and lentivirally transduced to express either CD19 linked in cis to green fluorescent protein (GFP) (referred to as target cells) via self-cleaving P2A or BCMA linked in *cis* to BFP (referred to as control cells) and purified by flow cytometry assisted sorting. 5×10^3^ target cells, 5×10^3^ control cells and 5×10^4^ CD8^+^ T cells were incubated with either various dilutions of supernatants from genome-engineered cells or media containing various concentrations of recombinant anti-CD19 x anti-CD3 bispecific (Invivogen, bimab-hcd19cd3) for 48 hours (Figure 1F-H & 3G-I). For the peripheral blood mononuclear cell (PBMC) killing assay, 2×10^5^ PBMCs and 4×10^4^ autologous CD8^+^ T cells were incubated with either supernatants from engineered PCs or media containing recombinant anti-CD19 x anti-CD3 bispecific (Invivogen, bimab-hcd19cd3) or anti-CD33 x anti-CD3 bispecific (AMG 330) for 48 hours (Figure 2H-K). For leukemia cell line killing assay 5×10^3^ NALM-6 cells, 5e3 MOLM-14 cells and 5×10^4^ CD8^+^ were incubated with either supernatants from engineered PCs or media containing recombinant bispecifics (Figure 2D-F). For the self-killing assay, 2×10^5^ genome-engineered B cells were incubated with autologous T cells at various effector to target ratios and cultured for 24 hours (Figure 3BE-F). Each assay was performed in 200uL in duplicate in 96 wells with RPMI-1640 supplemented with 10% FBS as the base media at 37°C and 5% CO_2_. At the end of each assay, cells from duplicate wells were pooled, washed with PBS, stained according to Table S2, and analyzed by flow cytometry.

### *In vivo* assessment of ePCs against human B-cell malignancies

All animal studies were performed according to AAALAC standards and were approved by the Seattle Children’s Research Institute (SCRI) Institutional Animal Care and Use Committee. NOD.Cg-*Prkdc^scid^ Il2rg^tm1Wjl^/*SzJ-c (NSG) mice were purchased from Jackson Laboratory and all mice were kept in a designated pathogen-free facility at SCRI. For the subcutaneous lymphoma flank model (Figure 4A-C), 2.5×10^5^ ePCs, 5×10^4^ autologous T cells, and 2.5×10^4^ luciferase transduced Raji cells (human Burkitt lymphoma cell line) were delivered subcutaneously to the right flank. For the disseminated leukemia experiments (Figure 5 and Figure 6), 2.5×10^6^-15×10^6^ GFP or bispecific-ePCs were injected intravenously into NSG. The following day (Figure 5), or the two days prior (Figure 6), mice received 1×10^5^ luciferase expressing B-ALL cells intravenously (model NL482B [Children’s Oncology Group unique specimen identifier PALJDL]). For Figure 5, the following day and three days later mice received 1×10^5^ or 1×10^6^ T cells administered retro-orbitally. For Figure 6, 2.5×10^6^ T cells were injected retro-orbitally the day following ePC engraftment, Tumor burden was monitored by bioluminescence imaging using IVIS Lumina S5 (Perkin Elmer) following subcutaneous injection of luciferin (75-150 mg/kg). Peripheral blood was collected via submandibular bleed and processed to collect sera and quantify human T cell numbers. Mice were euthanized for harvesting of bone marrow and spleens that were processed via erythrocyte lysis (ACK lysis) and then immunophenotyped by flow cytometry to quantify human leukemia cell and plasma cell numbers (Table S2).

**Figure 4:**
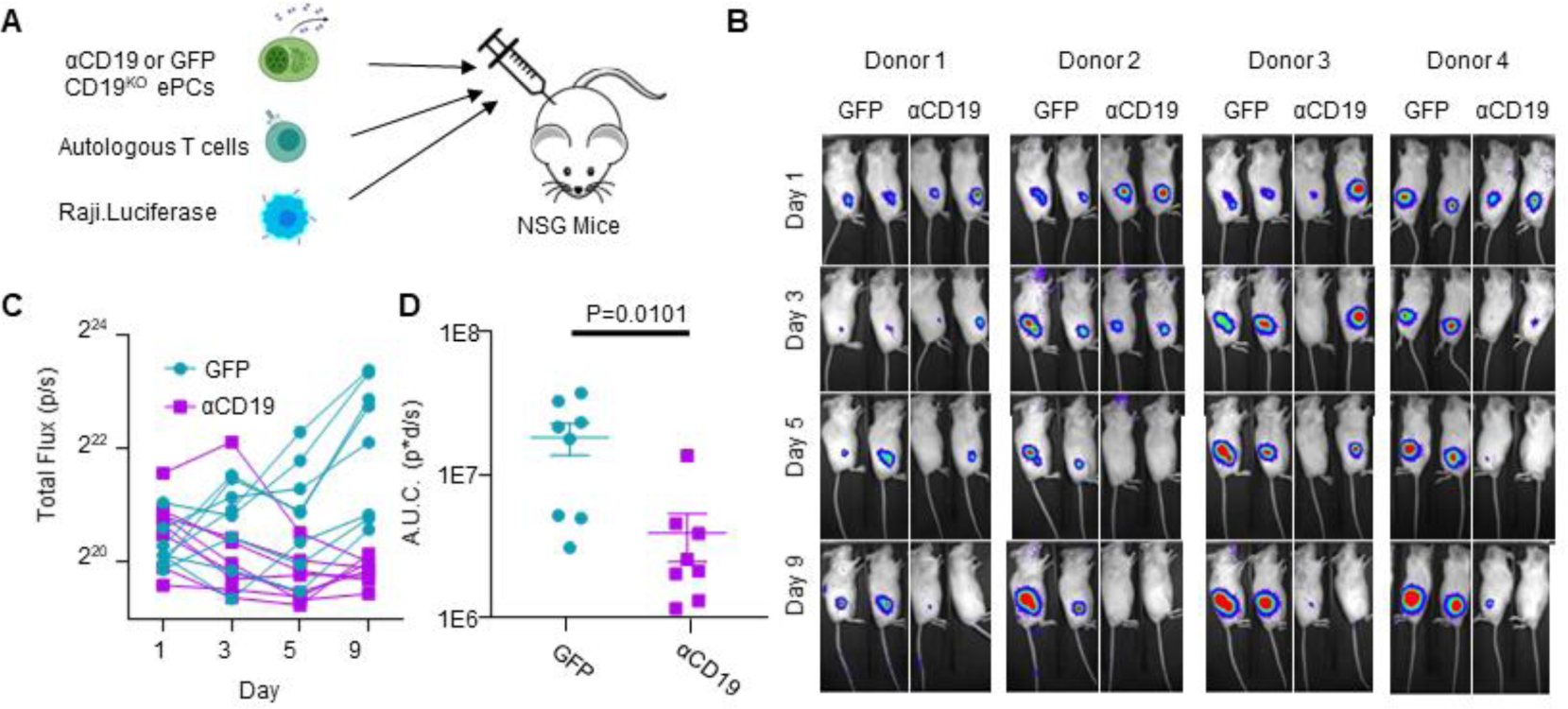
CD19^KO^ PCs engineered to secrete αCD19 bispecific have anti-lymphoma efficacy *in vivo*. **A)** Schematic showing an *in vivo* model for lymphoma growth. Briefly, GFP.CD19^KO^ or αCD19.GFP.CD19^KO^ ePCs, autologous T cells, and luciferase expressing Raji cells were injected subcutaneously into the right flank of immunodeficient NSG mice. **B)** Representative bioluminescence images were obtained via *in vivo* imaging (color scale; min:8×10^3^ max:1×10^5^). **C)** Bioluminescence was quantified from each mouse as total flux and graphed over time. **D)** Area under the curve analysis was conducted with baseline correction of 6×10^5^ flux. A-D) Data across 4 donors in two independent experiments with p-value calculated by unpaired student’s t test. Illustrations created in part with biorender.com.

**Figure 5:**
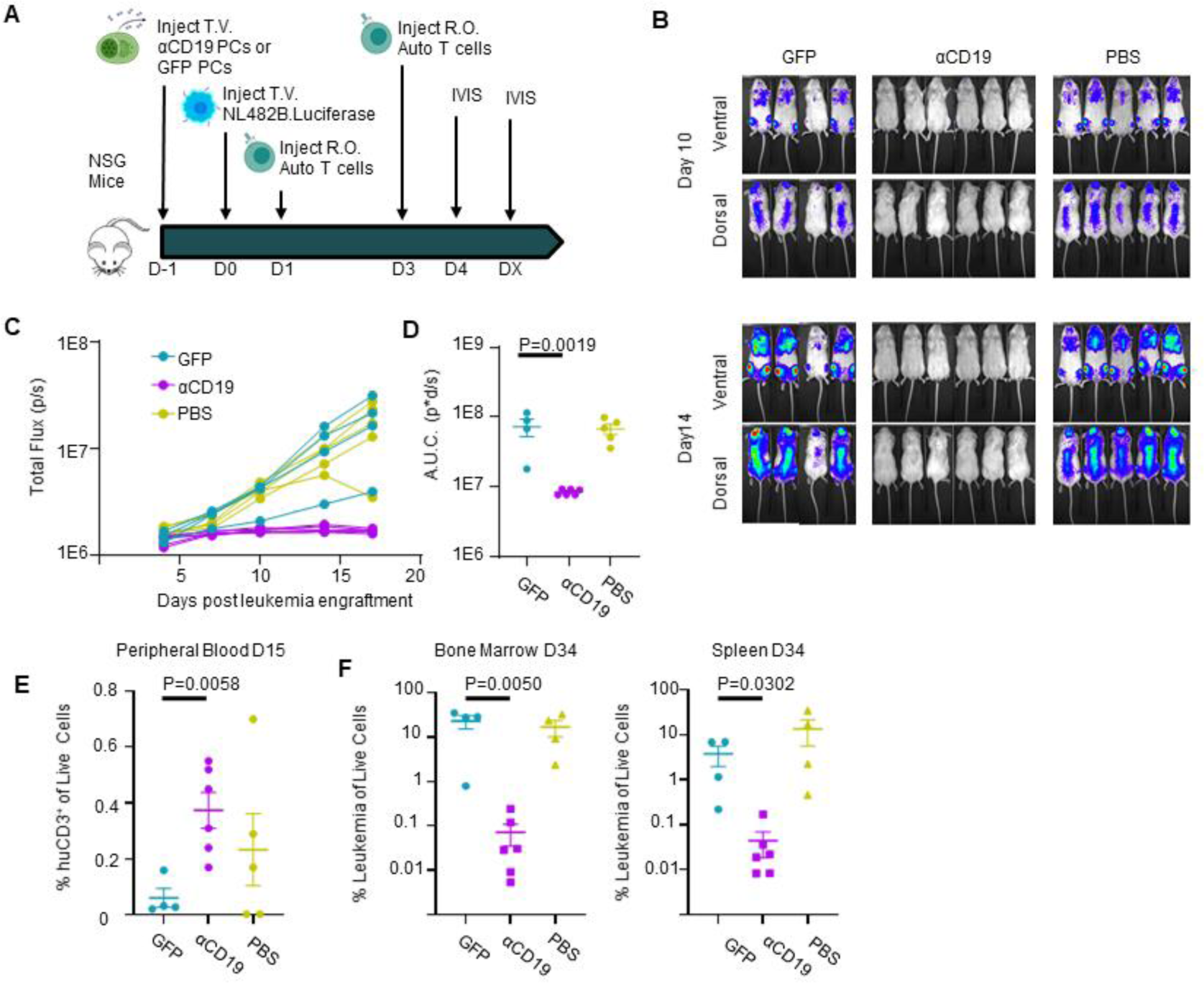
CD19^KO^ PCs engineered to secrete αCD19 bispecific can prevent leukemia engraftment. **A)** Schematic showing prophylactic treatment of a patient-derived xenograft model of high-risk ALL. Either GFP.CD19^KO^ or αCD19.GFP.CD19^KO^ ePCs were injected intravenously into immunodeficient NSG mice. 24 hours later, luciferase-labeled patient-derived NL482B [Children’s Oncology Group unique specimen identifier PALJDL] cells were administered. Finally, we delivered T cells syngeneic to the ePCs in two doses by retro-orbital injection. **B)** Bioluminescent images showing dissemination of the luciferase-expressing leukemia cells (color scale; min:8×10^3^ max:1×10^5^). **C)** Leukemia growth was quantified via total bioluminescent flux at the indicated time points. **D)** Area under the curve analysis was conducted with baseline correction 1×10^6^ flux. E) Peripheral blood flow analysis showing the percent of CD3+ cells of singlet live cells is elevated in the αCD19 cohort. Mice were euthanized 34 days after leukemia engraftment and tissues were stained and analyzed by flow. **F)** The percent CD19^+^ of live CD45^+^ singlet cells shows suppression of leukemic cells in bone and spleens of the αCD19 ePC cohort. A-D) Data from one donor with p-values calculated by one-way unpaired ANOVA with Šídák’s posttest (D) and unpaired student’s T tests between GFP and αCD19 cohorts (E-F). Illustrations created in part with biorender.com.

**Figure 6:**
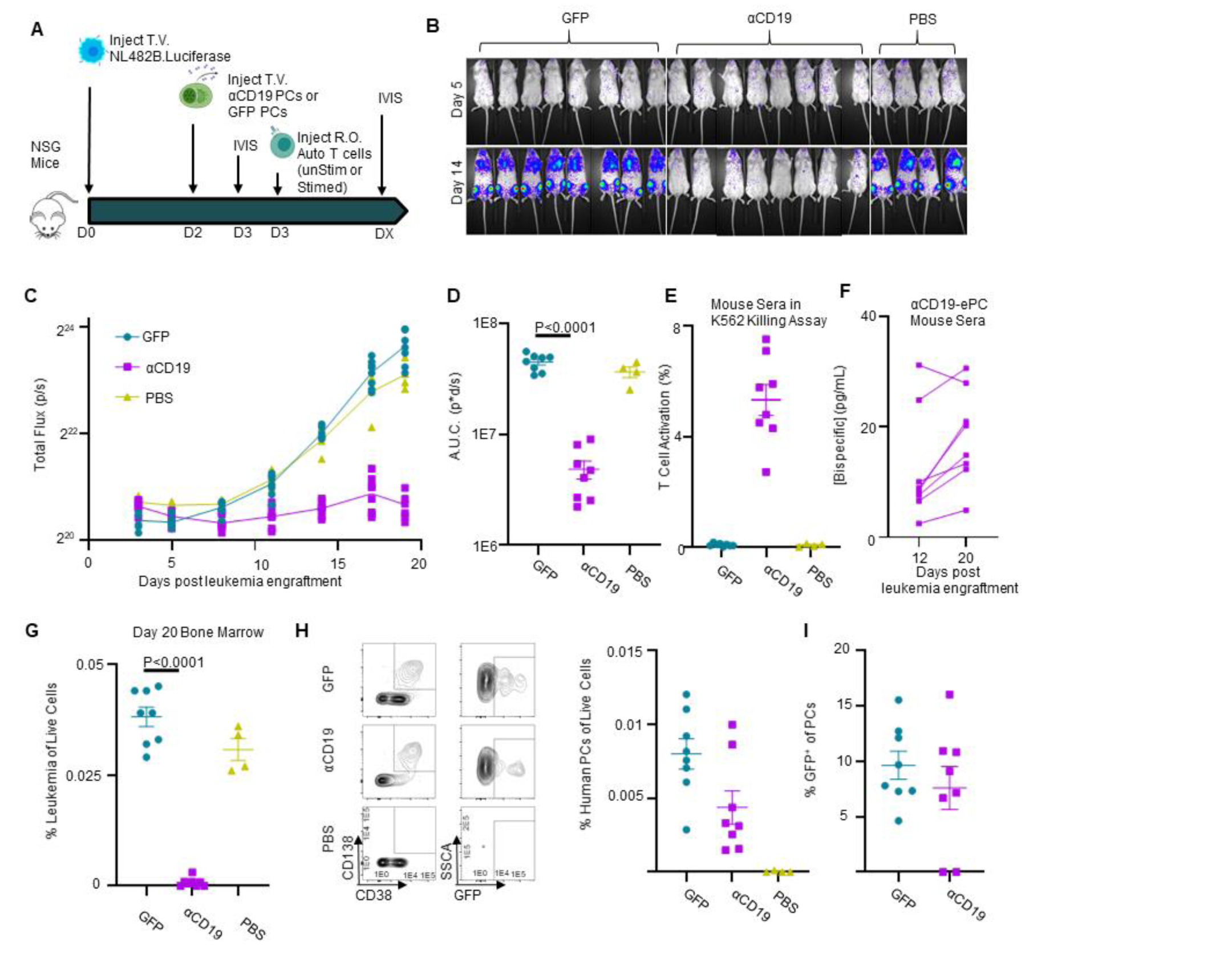
αCD19 bispecific secreting ePCs can persist in bone marrow and treat established leukemia. **A)** Schematic showing therapeutic treatment of a patient-derived xenograft model of high-risk ALL. Luciferase-labeled patient-derived NL482B [Children’s Oncology Group unique specimen identifier PALJDL] cells were administered intravenously. After 48 hours, either GFP.CD19^KO^ or αCD19.GFP.CD19^KO^ ePCs were injected intravenously into immunodeficient NSG mice. 24 hours later, we delivered T cells syngeneic to the ePCs via retro-orbital injection. **B)** Bioluminescent images showing dissemination of the luciferase-expressing leukemia cells (color scale; min:5×10e^3^ max:5×10e^4^). **C)** Leukemia growth was quantified via total bioluminescent flux at the indicated time points. **D)** Area under the curve analysis was conducted with baseline correction 1.25×10^6^ flux. Peripheral blood sera from mice at day 12 and day 20 was collected and used in the previously described K562 Killing assay. **E)** T cell action (%CD69^+^CD137^+^) caused by sera from mice twenty days post tumor engraftment is shown. **F)** Concentration of αCD19 bispecific in the mouse seras were interpolated from a standards curv*e.* Twenty days after tumor engraftment, bone marrow cells were harvested, stained, and analyzed by flow cytometry. **G)** The percent of tumor (huCD19^+^huCD45^+^moCD45^-^) of live cells was quantified. H) Representative flow plots of human cells show plasma cells present in the bone marrow of mice that received ePCs. The percent of plasma cells (huCD38^+^huCD45^+^moCD45-huCD138^+^) of live cells was calculated. **I)** The percentage of plasma cells that were GFP^+^ was quantified and plotted. Data from one donor with p-values calculated by one-way unpaired ANOVA with Šídák’s posttest. Illustrations created in part with biorender.com.

### Statistical analysis

Statistical analyses using parametric tests were performed using Prism 7 (GraphPad, San Diego, CA) as described in figure legends.

### Data sharing statement

For original data, reagents and protocols please contact.

## Results

### Primary human B cells engineered by HDR-based gene editing secrete functional bispecifics

To integrate a bispecific gene expression cassette into B cells, we adapted AAV-based HDR that we have used for delivery of transgenes in B cells at the safe harbor gene *CCR5*^13^. We designed *CCR5*-targeted HDR templates for delivery of Blue Fluorescent Protein (BFP) alone (as control) or an anti-CD19 x anti-CD3 bispecific cis-linked with GFP (heretofore referred to as a αCD19). We initiated gene editing by transfecting the activated human peripheral B cells with Cas9 ribonucleoprotein complexes (RNPs) containing guide RNAs targeting sequence within *CCR5*, and subsequently transduced with rAAV6 HDR donor vector (Figure 1A & S1A). We found that HDR integration rates were slightly lower with the vectors containing the αCD19 bispecifics when evaluated by digital droplet PCR (ddPCR; Figure S1B Figure 1B). However, despite similar integration rates the proportion of cells expressing the fluorescent reporter was substantially diminished in cells edited using the bispecific design, resulting in a significant drop in the ratio of fluorescent reporter marking to integration rate (Figure 1C-D).

We hypothesized that αCD19 bispecific expression could be increased by targeting transgene integration to loci that are natively expressed in B cells or plasma cells. Therefore, we built three additional AAV-based repair template designs for delivery of transgene cassettes to the highly expressed B cell loci *IGHG1*, *JCHAIN,* and a region proximal to an heavy chain enhancer, Eμ^18^ (repair arms and sgRNA were previously described in ^18^; overall schematic for all vectors, Figure 1E). Although the same ubiquitous viral-derived promoter (MND^33^) was used at all loci, we observed variable increases in GFP^+^ percentage and GFP mean fluorescent intensity in B cells following delivery to the antibody-associated loci, relative to that at *CCR5* (Figure S1C-D). While integration was detected at all loci, we observed significant increases in the mean fluorescent intensity of the cis-linked GFP at the antibody loci relative to *CCR5* (Figure 1F-G).

To initially assess the functionality of the B cell-produced αCD19 bispecific, we developed a fluorescent reporter-based *in vitro* killing assay using a K562 cell line that stably expresses CD19, a reference K562 cell line, CD8+ T cells incubated with either recombinant αCD19 bispecific or with supernatants from engineered cells (Figure H). After 48 hours, flow cytometry was used to quantify T cell activation (percent CD69^+^CD137^+^) and specific lysis of CD19^+^ target cells (Figure S2A). Recombinant αCD19 bispecific elicited dose and time-dependent increases in T cell activation and CD19-specific lysis (Figure S2B-C). Supernatants from B cells engineered to express the αCD19 bispecific induced T cell activation and specific lysis, whereas supernatants from GFP-engineered B cells did not (Figure 1I-J). We generated standard curves using recombinant bispecific T cell activation data to quantify the αCD19 bispecific concentrations in the supernatants derived from engineered B cells (Figure 1K). These findings indicate that primary human B cells can be engineered at various loci to express and secrete a functional αCD19 bispecific.

### Bispecific-ePCs exhibit *in vitro* activity against common leukemia target antigens

We next asked whether *ex vivo-*differentiated human ePCs could produce bispecifics that specifically target primary human hematopoietic cells expressing physiological levels of candidate leukemia antigens (including CD19 or CD33) within a heterogeneous cell population. We built an Eμ locus-directed bispecific vector for delivery of an anti-CD33 x anti-CD3 bispecific (heretofore referred to as a αCD33 bispecific; Figure 2A)^8^. We introduced each Eμ locus-directed bispecific or GFP alone control into B cells using HDR-based editing and differentiated the edited population into ePCs as previously described (Figure S1A).^13,32^ Following editing and differentiation, we observed detectable transgene expression with all vectors and donors (Figure 2B-C). Although we observed donor-dependent differences in relative expression of the plasma cell differentiation markers CD38 and CD138, introduction of the bispecifics did not impact differentiation into plasma cells (defined as CD38^++^ CD138^+^; Figure S3A-C), demonstrating that human plasma cells can be engineered to express bispecifics.

To investigate the functionality of bispecifics secreted by the ePCs to target physiological levels of antigen, we evaluated αCD19 and αCD33 ePC supernatants using two assays of heterologous cell populations. First, we applied recombinant bispecific to a PBMC killing assay wherein effector CD8^+^ T cells were co-cultured with autologous PBMCs that contained B cell and myeloid cell subpopulations expressing endogenous levels of CD19 and CD33 respectively (Figure 2D). As expected, recombinant αCD19 bispecific elicited a dose-dependent decrease in IgM^+^ B cells (Figure S4B), whereas recombinant αCD33 bispecific elicited a dose-dependent decrease in CD14^+^ CD33^+^ monocytes (Figure S4B). Supernatants from both αCD19- and αCD33-ePCs elicited higher T cell activation relative to that from control GFP-ePCs (Figure 2E). Furthermore, supernatants from αCD19-ePCs specifically lysed IgM^+^ B cells (Figure 2F), while supernatants from αCD33-ePCs specifically lysed CD33^+^ CD14^+^ monocytes (Figure 2G). Secondly, we evaluated ePC supernatants in a leukemia cell line killing assay wherein a CD19^+^ precursor B-ALL cell line (NALM-6), a CD33^+^ AML cell line (MOLM-14) and CD8^+^ effector T cells were co-cultured for 48 hours (Figure 2D). Increasing concentrations of recombinant αCD19 bispecific led to increased lysis of NALM-6 cells whereas increasing concentrations of recombinant αCD33 bispecific led to increased lysis of MOLM-14 cells (Figure S5A-B). T cells showed upregulation of activation markers CD69^+^ and CD137^+^ when cultured with supernatants from ePCs producing either bispecific relative to supernatants from control GFP-ePCs (Figure 2H). Supernatants from αCD19-ePCs specifically lysed NALM-6 cells (Figure 2I), whereas supernatants from αCD33-ePCs cells specifically lysed MOLM-14 cells (Figure 2J). Together, these data show that ePCs secreting αCD19 or αCD33 bispecifics elicit specific T cell killing of PBMC subsets or leukemia cell lines expressing physiological levels of CD19 or CD33 respectively.

### CD19^KO^ ePCs are protected from self-targeting and exhibit increased αCD19 bispecific secretion

CAR T cells engineered to recognize T-cell antigens can kill other CAR T cells within the same cell product, resulting in diminished anticancer activity^34,35^. We hypothesized that upon T cell encounter, αCD19-ePCs could similarly elicit self-targeting (ie fratricide) due to their surface CD19 expression (Figure S6). To evaluate the degree of self-targeting, we incubated Eμ locus-directed GFP-ePCs or αCD19-ePCs with autologous T cells (Figure 3A). Addition of autologous T cells lead to a progressive decline in the proportion of αCD19-ePCs but not GFP-ePCs (Figure 3B; gating Figure S7), implying that CD19 self-targeting likely impacts αCD19-secreting ePCs. Based on these observations, we predicted that elimination of CD19 would prevent αCD19 bispecific-elicited self-targeting.

To knockout CD19, we co-delivered RNPs targeting *CD19* with the Eμ locus-directed αCD19 bispecific editing reagents. The addition of the CD19-targeting RNPs resulted in >85% reduction in the proportion of CD19^+^ PCs (Figure 3C-D). CD19 knockout did not overtly impact differentiation of edited B cells into plasmablasts or PCs *in vitro* (Figure S8C). Upon challenging these CD19 knockout αCD19-ePCs with T cells, we observed no differences in GFP percentage with increased starting T cell numbers (Figure 3E). When comparing CD19^KO^ to CD19^wt^, only CD19^wt^ αCD19-ePCs exhibited a significant decrease in edited cells at the highest T cell dose (Figure 3F). These data suggest that CD19 knockout protects αCD19-ePCs from self-targeted death.

An additional challenge with CD19 surface expression is that the αCD19 bispecific produced from ePCs may bind to surface CD19 and limit the quantity that is released by the cells. Therefore, we predicted that knocking out CD19 would increase the level of αCD19 bispecific in supernatants. To assess free bispecific in the context of knocking out CD19, supernatants from αCD19 ePCs (co-engineered with or without *CD19* RNPs) were assessed using the K562 killing assay (Figure 1E). Supernatants from CD19^KO^ αCD19-ePCs resulted in higher T cell activation, and higher αCD19 bispecific concentrations and a trend towards higher specific lysis when compared to CD19^wt^ αCD19-ePCs (Figure 3G-I). Collectively these findings indicate that knocking out CD19 prevents self-targeting by T cells and boosts αCD19 bispecific levels.

Similar strategies could be employed for ePCs expressing biologics targeting additional B cell surface proteins. We individually knocked out the B cell surface markers *CD19*, *MS4A1* (CD20), *CD38* and *TNFRSF17 (*BCMA) to assess our ability to generate ePCs lacking these markers. Knock outs were confirmed by Inference of CRISPR Editing and by staining for surface expression (Supplemental 9A,C-D). None of the knockouts impacted cell expansion, viability, differentiation, or antibody secretion (Supplemental 9B-E). These data suggest that ePCs can be made that lack B cell associated antigens currently targeted in the clinic.

### αCD19 bispecific ePCs exhibit anti-tumor activity *in vivo*

To begin to test whether bispecific-secreting ePCs maintained function *in vivo*, we used a subcutaneous flank model of B cell lymphoma wherein luciferase-expressing lymphoma target cells, autologous T cells and ePCs were co-delivered to the flanks of immune deficient mice (Figure 4A). The lymphoma cells engrafted similarly in all groups (day 1 time point, Figure 4C-D). However, at later time points, tumor burden decreased in mice that received αCD19-ePCs relative to mice that received control GFP-ePCs (Figure 4B-D). In the αCD19-ePCs group, reductions in tumor size were below background luminescence levels in >50% of the mice within 5 days (Figure 4B-D). These findings demonstrate that αCD19-ePCs can promote robust local anti-tumor responses *in vivo*.

In some clinical settings, non-IgG-like αCD19 bispecifics are used to treat high-risk B-ALL patients as a bridge to hematopoietic stem cell transplantation.^36–41^ To potentially mimic this clinical scenario, we utilized a patient-derived, Philadelphia chromosome (PH)-like B-ALL xenograft model wherein CD19^KO^ GFP or αCD19-ePCs were adoptively transferred into immunodeficient mice and subsequently followed 1 day later with intravenous transfer of luciferase expressing PH-like B-ALL cells (NL482B; *IL7R* gain–of-function, *SH2B3* deletion).^42,43^ Effector T cells syngeneic with the ePCs were transferred retro-orbitally at 1 and 3 days after the B-ALL engraftment (Figure 5A). Control mice that received leukemia cells showed a steady increase in luciferase activity over time(Figure 5B-C). In contrast to control animals, mice that received αCD19-ePCs showed near complete leukemia control (Figure 5B-D). Consistent with these findings, the frequency of human T cells in the peripheral blood of the αCD19-ePC treated group trended higher, which is consistent with bispecific-driven T cell expansion *in vivo* (Figure 5E). Most importantly, the proportion of CD19^+^ leukemia cells was markedly reduced in both the spleen and bone marrow in the αCD19-ePC treated cohort upon sacrifice at 34 days post-leukemia cell transfer (Figure 5F). These findings demonstrate that bispecific-ePCs can limit in vivo growth and dissemination of a patient-derived leukemia in a B-ALL xenograft model.

Next, we tested the therapeutic potential of ePCs to treat established leukemia. Briefly, immunodeficient mice were intravenously engrafted with luciferase expressing PH-like B-ALL. After tumors were detectable by luciferase, CD19^KO^ GFP or αCD19-ePCs, and syngeneic T cells were adoptively transferred (Figure 6A). In contrast to control animals which exhibited increases in luciferase, mice that received αCD19-ePCs exhibited leukemia control (Figure 6B-D), exemplified by a slight decrease in luminescence between day 3 and day 8 post tumor engraftment (Figure 6C). Almost no leukemic cells were detectable in the bone marrow of the αCD19-ePC treated mice (Figure 6H). Upon quantifying bispecific levels in sera, we found that mice that received αCD19-ePCs had detectable signals in the T cell activation assay (Figure 6E). Furthermore, the concentration of bispecific in the sera of αCD19-ePCs remained stable between days 12 and 20 (Figure 6F). Consistent with a stable source of bispecific, αCD19-ePCs plasma cells that expressed the cis-linked GFP reporter could be detected in the bone marrow of mice 18 days after receiving αCD19-ePCs, but not in control mice that did not receive ePCs (Figure 6H-I). Together these findings suggest that αCD19-ePCs stably engraft in the bone marrow where they secrete αCD19 bispecific at detectable levels that are sufficient to mediate and maintain leukemia clearance *in vivo*.

## Discussion

Engineered plasma cells comprise an emerging cell-based modality for high-level, sustained delivery of therapeutic proteins; herein, we report the novel use of ePCs to produce bispecific therapeutics. Using HDR-based editing, expression of two alternative clinical bispecifics, αCD19 and αCD33, was achieved across a range of candidate loci actively expressed in primary human B cells and PCs (Figure 1). B cells engineered to express bispecifics could be differentiated into PCs that mediated specific killing of primary human cells and leukemia cell lines expressing CD19 or CD33, respectively (Figure 2). Further, knockout of the target antigen, CD19, led to a significant increase in functional αCD19 bispecific concentrations and prevented ePC self-targeting (Figure3). Finally, we show that αCD19 ePCs were capable of directing a T cell dependent anti-leukemia response against a locally engrafted cell line and, most notably, controlling the expansion of patient-derived leukemia xenograft, partially mimicking bispecific treatment in patients with high-risk B-ALL (Figure 4-6).

Persistent on-target off-tumor toxicity to normal bystander B cells is common in patients that respond to CD19-targeted chimeric antigen receptor (CAR) T cell therapy.^44–46^ Similarly, ePC therapies targeting lymphoid malignancies have the potential to cause B cell aplasia, hypogammaglobulinemia and long-term dysfunction of the immune system. These treatment related sequelae may last beyond the desired treatment window given that the ePCs persisted for at leasts 18 days and that the phenotype of bispecific ePCs-engineered cells described in this study (CD38^++^ CD138^+^) resembles the phenotype of long-lived PCs isolated from human bone marrow^47^. ePCs engineered using similar methods can persist in humanized mice >1 year^14^. Antibiotics, intravenous immunoglobulin replacement therapy as well as vaccinations effectively manage hypogammaglobulinemia and recurrent infections seen in CD19-CAR treated patients^48,49^, and may be effective for patients treated with αCD19 ePCs. To further mitigate on-target/off-tumor toxicity, ePCs could be engineered with a kill switch such as the clinically validated inducible caspase^50–52^ suicide gene system.

A potential barrier for use of ePCs for treatment of leukemia, and possibly other lymphoid malignancies, is that ePCs^14,15,18^ retain expression of surface markers targeted by many biologics (*eg.* CD19, CD20, CD38, and BCMA), which could result in self-targeting of the ePC. Consistent with this concept, we demonstrate that αCD19-ePCs express endogenous surface CD19 and are self-targeted in the presence of T cells. This phenomenon parallels similar findings in chimeric antigen receptor T cells engineered to target T cell-associated antigens (ie fratricide)^53–55^. As CD19 is not critical for PC function^44,56^, and is downregulated in long-lived PCs^47,56,57^, our engineering strategy for dual CD19 knockout and expression of αCD19 at Eμ locus is unlikely to impact the ePC function or longevity. An alternative strategy to also achieve this goal would be to engineer only at the *CD19* locus. Our data suggest that ePCs could be engineered to utilize a range of candidate bispecifics or monoclonal antibody-based therapeutics targeting B cell expressed tumor targets. Like *CD19*, knockout or depletion of the lymphoma and myeloma targets *MS4A1* (also known as CD20)^58–61^, and *CD38*^62,63^ in plasma cells does not acutely impact durable antibody titers, a corollary of their longevity and secretory capacity. Knockouts of *MS4A1* and *CD38* did not impair our ability to generate ePCs. Thus, our findings imply that similar strategies could be used to generate ePCs expressing biologics in use for chronic lymphocytic lymphoma (CD20; glofitamab^64^), non-Hodgkin’s lymphomas (CD20; rituximab^65^, odronextamab^66^, mosunetuzumab^67^) and multiple myeloma (CD38; daratumomab^68^, Bi38^69^). In contrast, knockout of *TNFRSF17* (also known as BCMA) in mice decreases PC survival and eliminates the antibody response^70^; hence, knockout of *TNFRSF17* would likely hamper ePC longevity and/or function.

Our findings suggest that ePCs may provide benefits for delivery of protein therapeutics beyond delivery of bispecifics as studied here. Therapeutic protein biologics were the second most approved drugs from 2009 to 2017^71^ and many suffer from suboptimal half-lives exemplified by blinatumomab^7^. Because of poor pharmacokinetics, many biologics used in chronic diseases require frequent (up to daily) and, in some cases, life-long dosing. Examples, include treatments for enzyme replacement (agalsidase beta; half-life of 56 to 76 minutes^72^, factor IX; 18 to 40 hours^73^, laronidase; 1.5 to 3.6 hours^74^), chronic autoimmune disorders (infliximab; 9.5 days^75^ etanercept; 80 hours^76^), diabetes (liraglutide; 13 hours^77^) and human immunodeficiency virus (enfuvirtide 3.4 hours^78^). The potential for ePCs to persist long term^14^ and produce robust levels of exogenous protein could be a key to unlocking the therapeutic potential of biologics or therapeutic peptides that lack efficacy due to poor pharmacokinetics.

In summary, these findings demonstrate the potential for human ePCs to mediate anti-leukemia responses and marks a key step in the realization of ePC as therapies to treat cancer, auto-immune disorders and protein deficiency disorders. Further studies in humanized mice and non-human primates are warranted to fully understand the activity, longevity, and tissue localization of ePCs.

## Acknowledgements

The authors graciously thank Andee Ott for bispecific protein purification assistance, Gene Hess for technical flow cytometry assistance, the Office of Animal Care for technical husbandry training and assistance, the SCRI viral core for making AAV6 virus and Ragan Pitner for general scientific writing support.

This research was supported by National Institutes of Health (NIH) grants F30 5F30AI164574-02 (T.F.H.; National Institute of Allergy and Infectious Disease [NIAID]). This work was also supported in part by the Seattle Children’s Research Institute (SCRI) Program for Cell and Gene Therapy (PCGT), the Children’s Guild Association Endowed Chair in Pediatric Immunology (to D.J.R.), and the Hansen Investigator in Pediatric Innovation Endowment (to D.J.R.). S.K.T. is a Scholar of the Leukemia & Lymphoma Society and holds the Joshua Kahan Endowed Chair in Pediatric Leukemia Research at the Children’s Hospital of Philadelphia. The work was also supported by the National Cancer Institute under activity number 5R01CA201135 (to R.G.J.). Finally, the work was supported by a research agreement with Be Biopharma. These funding sources had no influence on the design of the study; collection, management, analysis, and the interpretation of the data; preparation, review, or approval of the manuscript; and the decision to submit the manuscript for publication.

## Authorship Contributions

Contributions: K.R.T. and S.K.T. provided critical reagents and experimental guidance; T.F.H., P.N., and G.A., conducted experiments and acquired data; T.F.H., P.N., and R.G.J., analyzed data; T.F.H, R.G.J., and D.J.R. designed the study; T.F.H. and R.G.J. wrote the manuscript; T.FH, G.A., S.K.T., R.G.J. and D.J.R. edited the manuscript.

## References

1. Mullard A. FDA approves first CAR T therapy. Nat Rev Drug Discov. 2017;16(10):669.

2. Waldman AD, Fritz JM, Lenardo MJ. A guide to cancer immunotherapy: from T cell basic science to clinical practice. Nat Rev Immunol. 2020;20(11):651–668.

3. Siegel RL, Miller KD, Fuchs HE, Jemal A. Cancer Statistics, 2021. CA Cancer J Clin. 2021;71(1):7–33.

4. Kantarjian H, Stein A, Gökbuget N, et al. Blinatumomab versus Chemotherapy for Advanced Acute Lymphoblastic Leukemia. N Engl J Med. 2017;376(9):836–847.

5. Przepiorka D, Ko CW, Deisseroth A, et al. FDA Approval: Blinatumomab. Clin Cancer Res. 2015;21(18):4035–4039.

6. Halford Z, Coalter C, Gresham V, Brown T. A Systematic Review of Blinatumomab in the Treatment of Acute Lymphoblastic Leukemia: Engaging an Old Problem With New Solutions. Ann Pharmacother. 2021;55(10):1236–1253.

7. Zhu M, Wu B, Brandl C, et al. Blinatumomab, a Bispecific T-cell Engager (BiTE(®)) for CD-19 Targeted Cancer Immunotherapy: Clinical Pharmacology and Its Implications. Clin Pharmacokinet. 2016;55(10):1271–1288.

8. Ravandi F, Stein AS, Kantarjian HM, et al. A Phase 1 First-in-Human Study of AMG 330, an Anti-CD33 Bispecific T-Cell Engager (BiTE®) Antibody Construct, in Relapsed/Refractory Acute Myeloid Leukemia (R/R AML). Blood. 2018;132(Supplement 1):25–25.

9. Uy GL, Aldoss I, Foster MC, et al. Flotetuzumab as salvage immunotherapy for refractory acute myeloid leukemia. Blood. 2021;137(6):751–762.

10. Toksvang LN, Lee SHR, Yang JJ, Schmiegelow K. Maintenance therapy for acute lymphoblastic leukemia: basic science and clinical translations. Leukemia. 2022;36(7):1749–1758.

11. Apostolidou E, Lachowiez C, Juneja HS, et al. Clinical Outcomes of Patients With Newly Diagnosed Acute Lymphoblastic Leukemia in a County Hospital System. Clin Lymphoma Myeloma Leuk. 2021;21(11):e895–e902.

12. Strohl WR. Fusion Proteins for Half-Life Extension of Biologics as a Strategy to Make Biobetters. BioDrugs. 2015;29(4):215–239.

13. Hung KL, Meitlis I, Hale M, et al. Engineering Protein-Secreting Plasma Cells by Homology-Directed Repair in Primary Human B Cells. Mol Ther. 2018;26(2):456–467.

14. Cheng RYH, Hung KL, Zhang T, et al. Ex vivo engineered human plasma cells exhibit robust protein secretion and long-term engraftment in vivo. Nat Commun. 2022;13(1):6110.

15. Luo B, Zhan Y, Luo M, et al. Engineering of α-PD-1 antibody-expressing long-lived plasma cells by CRISPR/Cas9-mediated targeted gene integration. Cell Death Dis. 2020;11(11):973.

16. Page A, Laurent E, Nègre D, et al. Efficient adoptive transfer of autologous modified B cells: a new humanized platform mouse model for testing B cells reprogramming therapies. Cancer Immunol Immunother. 2022;71(7):1771–1775.

17. Levy C, Fusil F, Amirache F, et al. Baboon envelope pseudotyped lentiviral vectors efficiently transduce human B cells and allow active factor IX B cell secretion in vivo in NOD/SCIDγc-/-mice. J Thromb Haemost. 2016;14(12):2478–2492.

18. Moffett HF, Harms CK, Fitzpatrick KS, Tooley MR, Boonyaratanakornkit J, Taylor JJ. B cells engineered to express pathogen-specific antibodies protect against infection. Sci Immunol. 2019;4(35). doi:10.1126/sciimmunol.aax0644

19. Huang D, Tran JT, Olson A, et al. Vaccine Elicitation of HIV Broadly Neutralizing Antibodies from Engineered B cells.

20. Voss JE, Gonzalez-Martin A, Andrabi R, et al. Reprogramming the antigen specificity of B cells using genome-editing technologies. eLife. 2019;8.

21. Hartweger H, McGuire AT, Horning M, et al. HIV-specific humoral immune responses by CRISPR/Cas9-edited B cells. J Exp Med. 2019;216(6):1301–1310.

22. Nahmad AD, Raviv Y, Horovitz-Fried M, et al. Engineered B cells expressing an anti-HIV antibody enable memory retention, isotype switching and clonal expansion. Nat Commun. 2020;11(1):5851.

23. Page A, Hubert J, Fusil F, Cosset FL. Exploiting B Cell Transfer for Cancer Therapy: Engineered B Cells to Eradicate Tumors. Int J Mol Sci. 2021;22(18). doi:10.3390/ijms22189991

24. Manz RA, Thiel A, Radbruch A. Lifetime of plasma cells in the bone marrow. Nature. 1997;388(6638):133-134.

25. Amanna IJ, Carlson NE, Slifka MK. Duration of humoral immunity to common viral and vaccine antigens. N Engl J Med. 2007;357(19):1903–1915.

26. Eyer K, Doineau RCL, Castrillon CE, et al. Single-cell deep phenotyping of IgG-secreting cells for high-resolution immune monitoring. Nat Biotechnol. 2017;35(10):977–982.

27. Radbruch A, Muehlinghaus G, Luger EO, et al. Competence and competition: the challenge of becoming a long-lived plasma cell. Nat Rev Immunol. 2006;6(10):741–750.

28. Hibi T, Dosch HM. Limiting dilution analysis of the B cell compartment in human bone marrow. Eur J Immunol. 1986;16(2):139–145.

29. Marchand T, Pinho S. Leukemic Stem Cells: From Leukemic Niche Biology to Treatment Opportunities. Front Immunol. 2021;12:775128.

30. Benet Z, Jing Z, Fooksman DR. Plasma cell dynamics in the bone marrow niche. Cell Rep. 2021;34(6):108733.

31. Tasian SK, Bornhäuser M, Rutella S. Targeting Leukemia Stem Cells in the Bone Marrow Niche. Biomedicines. 2018;6(1). doi:10.3390/biomedicines6010022

32. Cheng RYH, de Rutte J, Ott AR, et al. SEC-seq: Association of molecular signatures with antibody secretion in thousands of single human plasma cells. bioRxiv. Published online August 26, 2022:2022.08.25.505190. doi:10.1101/2022.08.25.505190

33. Robbins PB, Yu XJ, Skelton DM, et al. Increased probability of expression from modified retroviral vectors in embryonal stem cells and embryonal carcinoma cells. J Virol. 1997;71(12):9466–9474.

34. Fleischer LC, Spencer HT, Raikar SS. Targeting T cell malignancies using CAR-based immunotherapy: challenges and potential solutions. J Hematol Oncol. 2019;12(1):141.

35. Gower M, Tikhonova AN. Avoiding fratricide: a T-ALL order. Blood. 2022;140(1):3–4.

36. Keating AK, Gossai N, Phillips CL, et al. Reducing minimal residual disease with blinatumomab prior to HCT for pediatric patients with acute lymphoblastic leukemia. Blood Adv. 2019;3(13):1926–1929.

37. Pawinska-Wasikowska K, Wieczorek A, Balwierz W, Bukowska-Strakova K, Surman M, Skoczen S. Blinatumomab as a Bridge Therapy for Hematopoietic Stem Cell Transplantation in Pediatric Refractory/Relapsed Acute Lymphoblastic Leukemia. Cancers . 2022;14(2). doi:10.3390/cancers14020458

38. Bargou R, Leo E, Zugmaier G, et al. Tumor regression in cancer patients by very low doses of a T cell-engaging antibody. Science. 2008;321(5891):974-977.

39. Mølhøj M, Crommer S, Brischwein K, et al. CD19-/CD3-bispecific antibody of the BiTE class is far superior to tandem diabody with respect to redirected tumor cell lysis. Mol Immunol. 2007;44(8):1935–1943.

40. Dreier T, Lorenczewski G, Brandl C, et al. Extremely potent, rapid and costimulation-independent cytotoxic T-cell response against lymphoma cells catalyzed by a single-chain bispecific antibody. Int J Cancer. 2002;100(6):690–697.

41. Dreier T, Baeuerle PA, Fichtner I, et al. T cell costimulus-independent and very efficacious inhibition of tumor growth in mice bearing subcutaneous or leukemic human B cell lymphoma xenografts by a CD19-/CD3-bispecific single-chain antibody construct. J Immunol. 2003;170(8):4397–4402.

42. Bride KL, Hu H, Tikhonova A, et al. Rational drug combinations with CDK4/6 inhibitors in acute lymphoblastic leukemia. Haematologica. 2022;107(8):1746–1757.

43. Maude SL, Tasian SK, Vincent T, et al. Targeting JAK1/2 and mTOR in murine xenograft models of Ph-like acute lymphoblastic leukemia. Blood. 2012;120(17):3510–3518.

44. Bhoj VG, Arhontoulis D, Wertheim G, et al. Persistence of long-lived plasma cells and humoral immunity in individuals responding to CD19-directed CAR T-cell therapy. Blood. 2016;128(3):360–370.

45. Maude SL, Frey N, Shaw PA, et al. Chimeric antigen receptor T cells for sustained remissions in leukemia. N Engl J Med. 2014;371(16):1507–1517.

46. Hill JA, Li D, Hay KA, et al. Infectious complications of CD19-targeted chimeric antigen receptor-modified T-cell immunotherapy. Blood. 2018;131(1):121–130.

47. Halliley JL, Tipton CM, Liesveld J, et al. Long-Lived Plasma Cells Are Contained within the CD19(-)CD38(hi)CD138(+) Subset in Human Bone Marrow. Immunity. 2015;43(1):132–145.

48. Topp M, Feuchtinger T. Management of Hypogammaglobulinaemia and B-Cell Aplasia. Springer; 2022.

49. Kampouri E, Walti CS, Gauthier J, Hill JA. Managing hypogammaglobulinemia in patients treated with CAR-T-cell therapy: key points for clinicians. Expert Rev Hematol. 2022;15(4):305–320.

50. Straathof KC, Pulè MA, Yotnda P, et al. An inducible caspase 9 safety switch for T-cell therapy. Blood. 2005;105(11):4247–4254.

51. Di Stasi A, Tey SK, Dotti G, et al. Inducible apoptosis as a safety switch for adoptive cell therapy. N Engl J Med. 2011;365(18):1673–1683.

52. Stavrou M, Philip B, Traynor-White C, et al. A Rapamycin-Activated Caspase 9-Based Suicide Gene. Mol Ther. 2018;26(5):1266–1276.

53. Cooper ML, Choi J, Staser K, et al. An “off-the-shelf” fratricide-resistant CAR-T for the treatment of T cell hematologic malignancies. Leukemia. 2018;32(9):1970–1983.

54. Mamonkin M, Rouce RH, Tashiro H, Brenner MK. A T-cell-directed chimeric antigen receptor for the selective treatment of T-cell malignancies. Blood. 2015;126(8):983–992.

55. Gomes-Silva D, Srinivasan M, Sharma S, et al. CD7-edited T cells expressing a CD7-specific CAR for the therapy of T-cell malignancies. Blood. 2017;130(3):285–296.

56. Mei HE, Wirries I, Frölich D, et al. A unique population of IgG-expressing plasma cells lacking CD19 is enriched in human bone marrow. Blood. 2015;125(11):1739–1748.

57. Arumugakani G, Stephenson SJ, Newton DJ, et al. Early Emergence of CD19-Negative Human Antibody-Secreting Cells at the Plasmablast to Plasma Cell Transition. J Immunol. 2017;198(12):4618–4628.

58. Hammarlund E, Thomas A, Amanna IJ, et al. Plasma cell survival in the absence of B cell memory. Nat Commun. 2017;8(1):1781.

59. Kuijpers TW, Bende RJ, Baars PA, et al. CD20 deficiency in humans results in impaired T cell-independent antibody responses. J Clin Invest. 2010;120(1):214–222.

60. Kozlova V, Ledererova A, Ladungova A, et al. CD20 is dispensable for B-cell receptor signaling but is required for proper actin polymerization, adhesion and migration of malignant B cells. PLoS One. 2020;15(3):e0229170.

61. Langley WA, Wieland A, Ahmed H, et al. Persistence of Virus-Specific Antibody after Depletion of Memory B Cells. J Virol. 2022;96(9):e0002622.

62. Hotchandani N, Fung H, Dulaimi E. CD38 Expression Loss in Multiple Myeloma Treated with Daratumumab. Am J Clin Pathol. 2016;146(suppl_1). doi:10.1093/ajcp/aqw151.007

63. Postigo J, Iglesias M, Cerezo-Wallis D, et al. Mice deficient in CD38 develop an attenuated form of collagen type II-induced arthritis. PLoS One. 2012;7(3):e33534.

64. Papazoglou D, Ysebaert L, Ioannou N, et al. S141: ELICITING ANTI-TUMOR T CELL ACTIVITY IN CHRONIC LYMPHOCYTIC LEUKEMIA WITH BISPECIFIC ANTIBODY-BASED COMBINATION THERAPY. HemaSphere. 2022;6:42.

65. Dotan E, Aggarwal C, Smith MR. Impact of Rituximab (Rituxan) on the Treatment of B-Cell Non-Hodgkin’s Lymphoma. P T. 2010;35(3):148–157.

66. Bannerji R, Arnason JE, Advani RH, et al. Odronextamab, a human CD20×CD3 bispecific antibody in patients with CD20-positive B-cell malignancies (ELM-1): results from the relapsed or refractory non-Hodgkin lymphoma cohort in a single-arm, multicentre, phase 1 trial. Lancet Haematol. 2022;9(5):e327–e339.

67. Budde LE, Assouline S, Sehn LH, et al. Single-Agent Mosunetuzumab Shows Durable Complete Responses in Patients With Relapsed or Refractory B-Cell Lymphomas: Phase I Dose-Escalation Study. J Clin Oncol. 2022;40(5):481–491.

68. Abdallah N, Kumar SK. Daratumumab in untreated newly diagnosed multiple myeloma. Ther Adv Hematol. 2019;10:2040620719894871.

69. Fayon M, Martinez-Cingolani C, Abecassis A, et al. Bi38-3 is a novel CD38/CD3 bispecific T-cell engager with low toxicity for the treatment of multiple myeloma. Haematologica. 2021;106(4):1193–1197.

70. O’Connor BP, Raman VS, Erickson LD, et al. BCMA is essential for the survival of long-lived bone marrow plasma cells. J Exp Med. 2004;199(1):91–98.

71. Batta A, Kalra BS, Khirasaria R. Trends in FDA drug approvals over last 2 decades: An observational study. J Family Med Prim Care. 2020;9(1):105–114.

72. Clarke JTR, West ML, Bultas J, Schiffmann R. The pharmacology of multiple regimens of agalsidase alfa enzyme replacement therapy for Fabry disease. Genet Med. 2007;9(8):504–509.

73. Chhabra A, Spurden D, Fogarty PF, et al. Real-world outcomes associated with standard half-life and extended half-life factor replacement products for treatment of haemophilia A and B. Blood Coagul Fibrinolysis. 2020;31(3):186–192.

74. Jameson E, Jones S, Remmington T. Enzyme replacement therapy with laronidase (Aldurazyme®) for treating mucopolysaccharidosis type I. Cochrane Database Syst Rev. 2019;6(6):CD009354.

75. Cornillie F, Shealy D, D’Haens G, et al. Infliximab induces potent anti-inflammatory and local immunomodulatory activity but no systemic immune suppression in patients with Crohn’s disease. Aliment Pharmacol Ther. 2001;15(4):463–473.

76. Nestorov I, Zitnik R, DeVries T, Nakanishi AM, Wang A, Banfield C. Pharmacokinetics of subcutaneously administered etanercept in subjects with psoriasis. Br J Clin Pharmacol. 2006;62(4):435–445.

77. Sisson EM. Liraglutide: clinical pharmacology and considerations for therapy. Pharmacotherapy. 2011;31(9):896–911.

78. Cheng S, Wang Y, Zhang Z, et al. Enfuvirtide-PEG conjugate: A potent HIV fusion inhibitor with improved pharmacokinetic properties. Eur J Med Chem. 2016;121:232–237.

