## Supplement for "Human plasma cells engineered to secrete bispecifics drive effective *in vivo* leukemia killing"

### Supplemental Materials

#### Supplemental Methods

##### Cell lines

K562, Raji, were obtained from the ATCC and cultured in RPMI 1640 supplemented with 10% fetal bovine serum. K562 were retroviral transduced with VSV-g pseudotyped virus containing CD19 in cis to GFP or BCMA (NP_001183) in cis to BFP and subsequently purified using fluorescence activated cell sorting (FACS) to generate target K562 $CD19^{+}$ $GFP^{+}$ and control reference K562 $BCMA^{+}$ $BFP^{+}$ cell lines. Raji were lentivirally transduced with firefly luciferase (AB261984.1) and enhanced green fluorescent protein (HM640279.1) and FACS-purified. NL428B cells expressing luciferase were provided by K. Thomas and S. Tasian. MOLM-14 cells lentiviral transduced to express GFP and luciferase were provided by S. Tasian.

##### Flow cytometry

Cells were stained according to staining panels in supplementary table 2. Flow cytometric analysis was performed on an LSR II flow cytometer (BD Biosciences) and events were analyzed using FlowJo software (Tree Star). Flow cytometry gating for fluorescent proteins, viability, and immunophenotyping can be found in supplementary figures.

#### Supplemental Figure Legends

##### Supplemental 1: High GFP editing efficiencies achieved at multiple loci

**A)** Schematic showing the protocols used for B cell expansion and differentiation. B cells were enriched from human PBMCs and cultured in activating media for two days. B cells were then engineered using homologous directed repair gene editing. Engineered cells were further expanded for 5 days. For figure 1 experiments only: engineered expanded B cells were harvested on day 7 for ddPCR, flow cytometry, and supernatants for killing assays. Cells were then cultured at 1E6/mL in cytokines driving differentiation towards PCs. **B)** Schematic showing the primer design for quantification of homology directed repair upon integration at the CCR5 locus. Integrated alleles and reference alleles were measured by ddPCR and used to calculate HDR allele frequency in Figure 1B. **C)** Gating strategy for determining the %GFP$^{+}$ and GFP MFI of edited B cells at Day 7. Flow data was first gated on lymphocytes by SSC-A vs FSC-A. Lymphocytes were then gated on live cells (AF350$^{-}$). Live lymphocytes were then gated for single cells and analyzed for GFP expression. **D)** %GFP$^{+}$ was calculated from singlet live cells and the mean fluorescent index was calculated from GFP$^{+}$ cells.

**Supplemental 2: Recombinant bispecific results in dose dependent T cell activation and CD19-specific lysis in a K562 killing assay**

Various concentrations of recombinant αCD19 bispecific were cultured with target (GFP$^{+}$:CD19$^{+}$) and control reference (BFP$^{+}$:CD19$^{-}$) K562 cells and CD8$^{+}$ T cells. Cells were harvested for analysis by flow cytometry after 48 hours. **A)** Gating strategy for flow cytometry of K562 killing assay. T cell activation is defined as the percent of live CD8$^{+}$ cells that are CD69$^{+}$ CD137$^{+}$. Specific lysis is calculated using the ratio of GFP$^{+}$:CD19$^{+}$ to BFP$^{+}$:CD19$^{-}$ cells normalized to the same ratio in the controls that received media only. **B)** Standard curves were interpolated for each experiment. Recombinant anti-CD19, anti-CD3 bispecific antibodies were added to the K562-CD8 T cell co-cultures. At the indicated concentration of bispecific, we quantified specific lysis and T cell activation as described in A). These standard curves were then used to back calculate the concentration of the supernatants from ePCs engineered to secrete the αCD19 bispecific in Fig 1J.

**Supplemental 3: Cells differentiated to day 10 maintain bispecific transgene expression and express phenotypic markers of plasma cells**

B cells were enriched from human PBMCs and cultured in activating media for two days. B cells were then engineered to express GFP or αCD19.T2A.GFP or αCD33.T2A.GFP transgenes. Engineered cells were further expanded for 5 days and then cultured at 1E6/mL in cytokines driving differentiation towards PCs for an additional 3 days. Engineered PCs analyzed for PC phenotype by flow cytometry, and supernatants were used for PBMC killing assays. **A)** Gating strategy for flow cytometry analysis of the PC phenotypes. **B)** Quantification of PC phenotype ($CD38^+$ $CD138^{+)}$ of singlet live lymphocytes in cells engineered to express the indicated bispecific antibodies. **C)** Representative flow plots and quantification of transgene positivity in bulk D10 cells or in the $CD38^+$ $CD138^+$ PC subsets.

**Supplemental 4: PBMC killing assay shows specific lysis and specific T cell activation in a dose dependent manner**

Human PBMCs and autologous $CD8^+$ T cells were co-cultured in the presence of either recombinant αCD19 bispecific or recombinant αCD33 bispecific for 48 hours. **A)** Cells were harvested and analyzed using the flow gating strategy shown. **B)** Cell frequencies were quantified and plotted by concentration of either αCD19 bispecific or αCD33 bispecific. Standards curves were interpolated by four parameter logistic based on T cell activation data.

**Supplemental 5: Bispecifics drive T cell dependent specific lysis in a leukemia cell line killing assay**

NALM-6 (B-ALL), MOLM-14 (AML) and autologous CD8+ T cells were co-cultured in the presence of either recombinant αCD19 bispecific or recombinant αCD33 bispecific for 48 hours. **A)** Cells were harvested and analyzed by flow cytometry using the gating strategy shown. **B)** NALM-6 ($IgM^+$) and MOLM-14 ($CD33^+$) cell frequencies were quantified and plotted as a function of the indicated bispecific concentrations. **C)** T cell activation ($CD69^+CD137^+$ of $CD8^+$) were quantified and plotted as a function of the indicated bispecific. Standard curves were interpolated by four parameter logistic based on T cell activation data.

**Supplemental 6: CD19 surface staining is lower on αCD19 engineered plasma cells**

Primary human B cells were enriched from PBMCs and cultured in activating media for two days and then engineered to express GFP or αCD19.T2A.GFP at the Eμ locus. Engineered cells were then expanded and subsequently differentiated into plasma cells over an additional 8 days. **A)** Gating strategy to analyze CD19 expression on engineered cells ($GFP^+$). **B)** Quantification of CD19 mean fluorescent intensity on engineered cells ($GFP^+$).

**Supplemental 7: Flow cytometry gating Strategy for B cell self-targeting assay**

Primary human B cells were enriched from PBMCs and cultured in activating media for two days and then engineered to express GFP or αCD19.T2A.GFP at the Eμ locus with or without CD19 knockout. Engineered cells were expanded for an additional 5 days and then incubated with autologous T cells. After 24hrs the percentage of edited B cells was determined by flow cytometry according to the gating strategy above.

**Supplemental 8: CD19 knockout does not affect differentiation of bispecific engineered B cells into plasma cells**

Primary human B cells were enriched from PBMCs and cultured in activating media for two days and then engineered to express GFP or αCD19.T2A.GFP at the Eμ locus with or without CD19 knockout. Edited cells were expanded for an additional 5 days. **A)** A trend towards lower viability was seen in cells that had CD19 knocked out. **B)** However, cell count remained high by D7. Engineered cells were further differentiated for 3 days into PCs. **C)** Cells were stained for surface markers and quantified as PCs ($CD38^{++}CD138^{+}$) and plasmablasts ($CD38^{++}CD138^{-}$).

**Supplemental 9: Engineered plasma cells secreting an αCD19 bispecific drive T cell expansion and CD19+ leukemia eradication *in vivo***

Peripheral blood was collected 15 days post leukemia engraftment. Spleens and bone marrow were collected 34 days post leukemia engraftment. Red blood cells were lysed, and remaining cells were stained for mouse and human surface markers. **A)** Gating strategy used for determining cell percentages from tissues. **B)** The percent of CD3+ cells of singlet live cells shows elevated T cells in αCD19 cohort. **C)** The percent $CD19^{+}$ of live $CD45^{+}$ singlet cells shows suppression of leukemic cells in the αCD19 cohort. P-values were calculated using a paired one-way ANOVA with Dunnett's posttest.

### Supplementary Figure 1

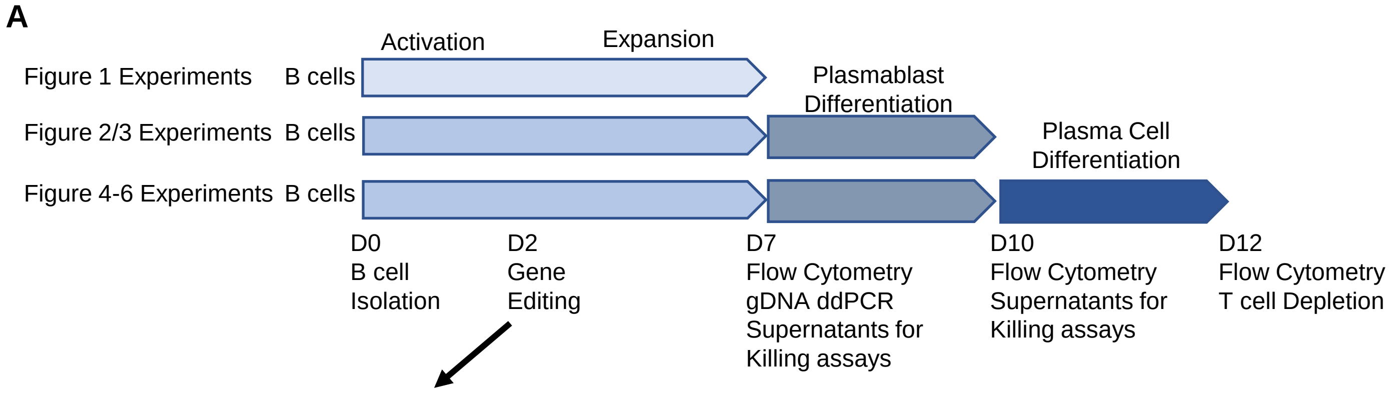


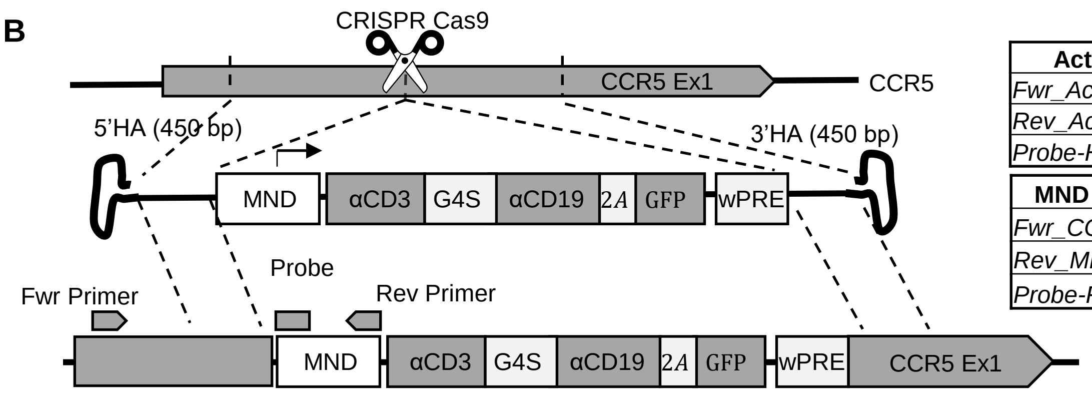


| ActB Ref | Sequence |
|---|---|
| *Fwr_ActB* | ACTCTGCAGGTTCTATTTGC |
| *Rev_ActB* | AATGATCTGAGGAGGGAAGG |
| *Probe-Hex* | ATCAAGGTGGGTGTCTTTCC |

| MND Target | Sequence |
|---|---|
| *Fwr_CCR5* | GGTATGATGCTTAGAACAGTGAT |
| *Rev_MND* | CCATATTCTGCTGTTCCAACT |
| *Probe-Fam* | CCTGGGCAACATAGTGTGAT |

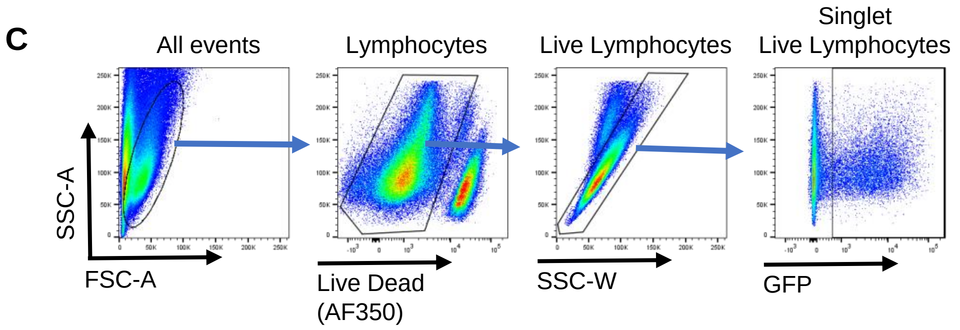


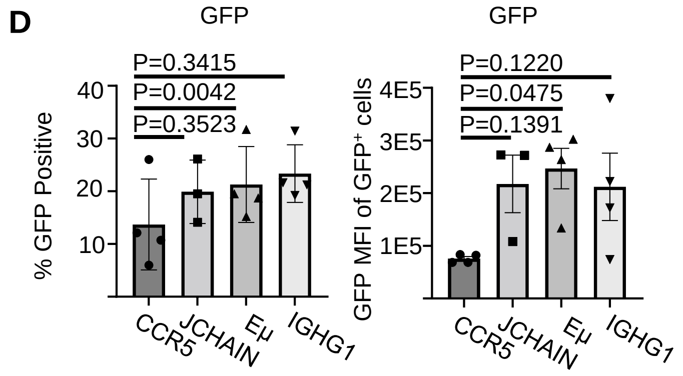

### Supplementary Figure 2

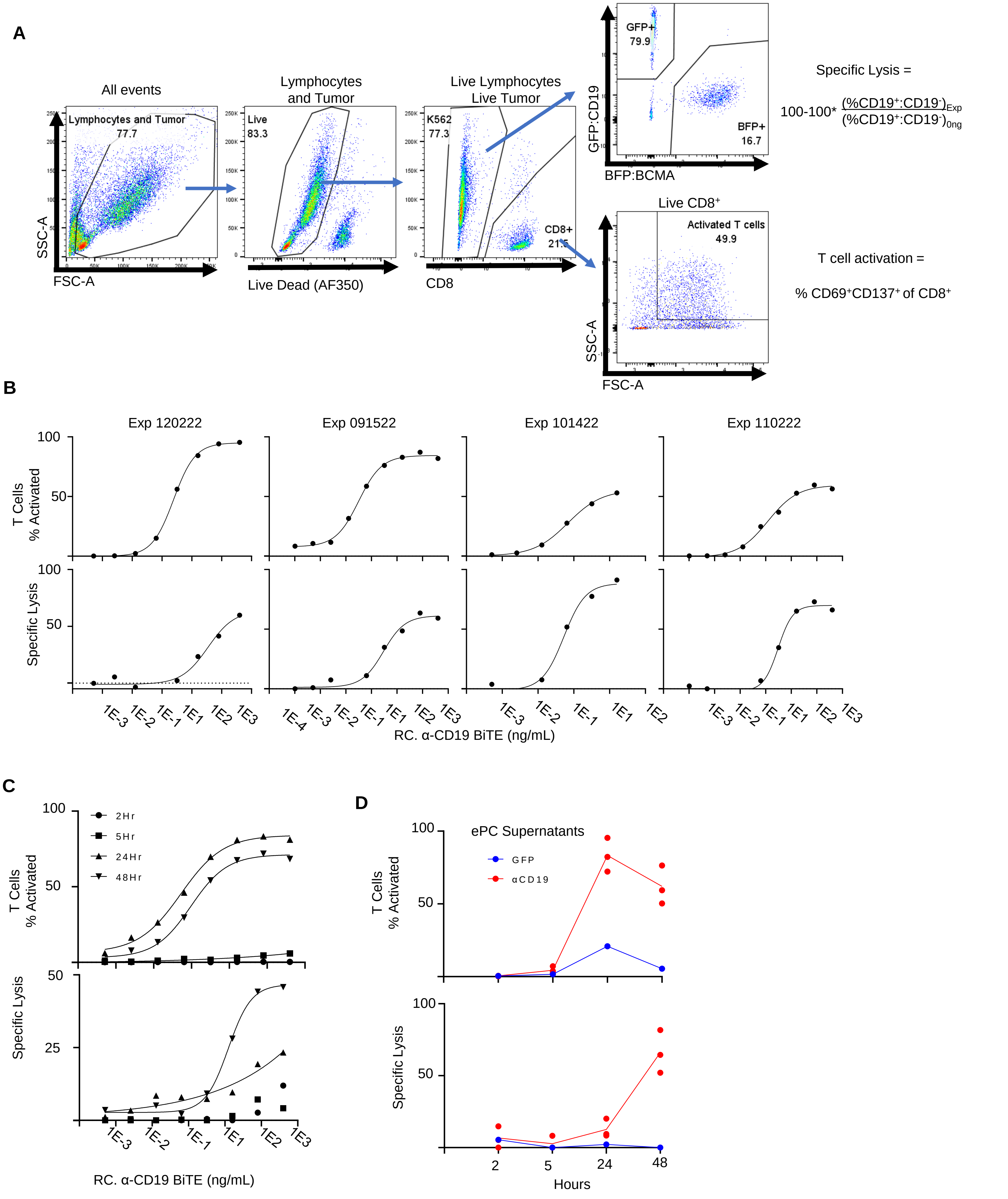

### Supplementary Figure 3

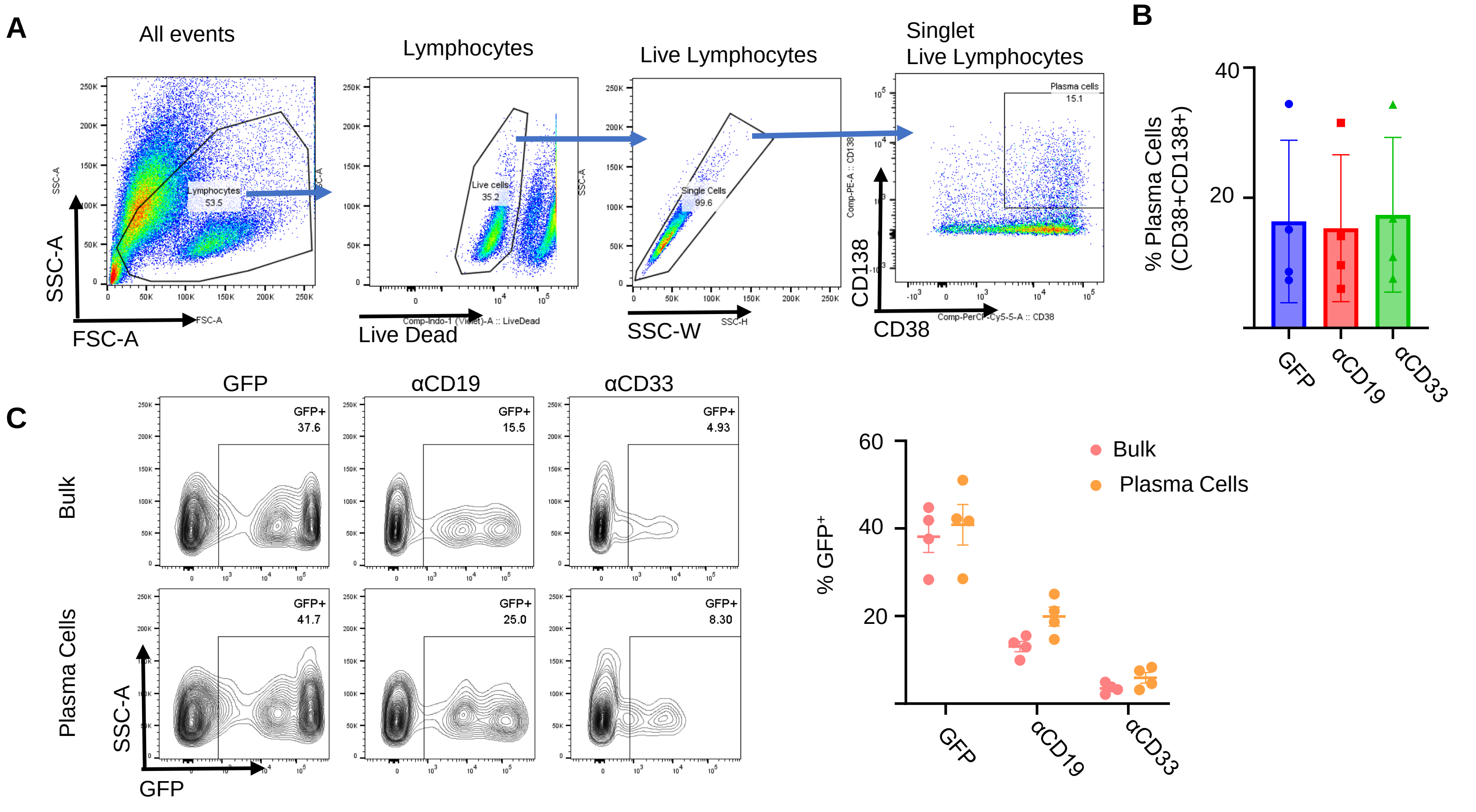


### Supplementary Figure 4

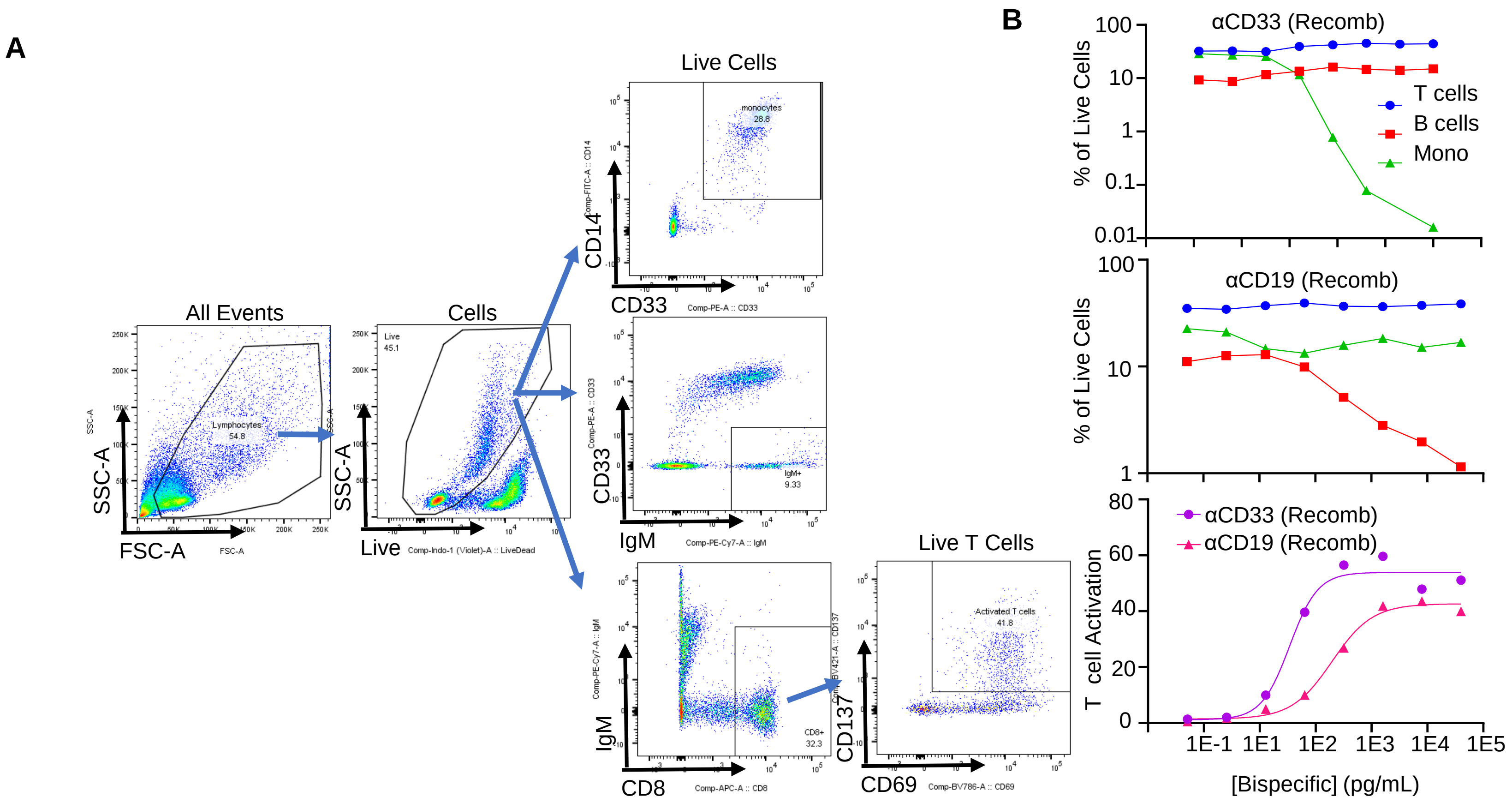

### Supplementary Figure 5

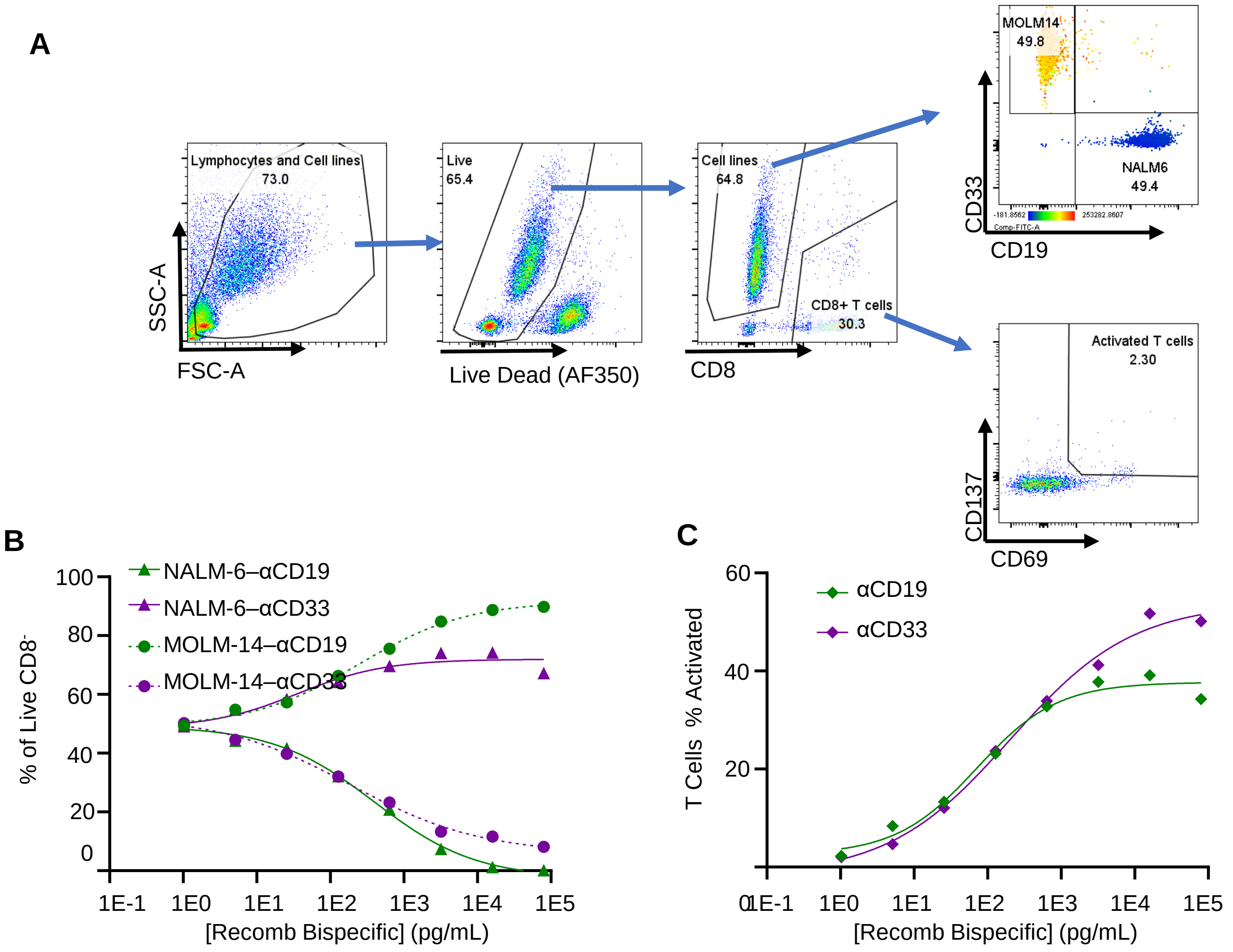

### Supplementary Figure 6

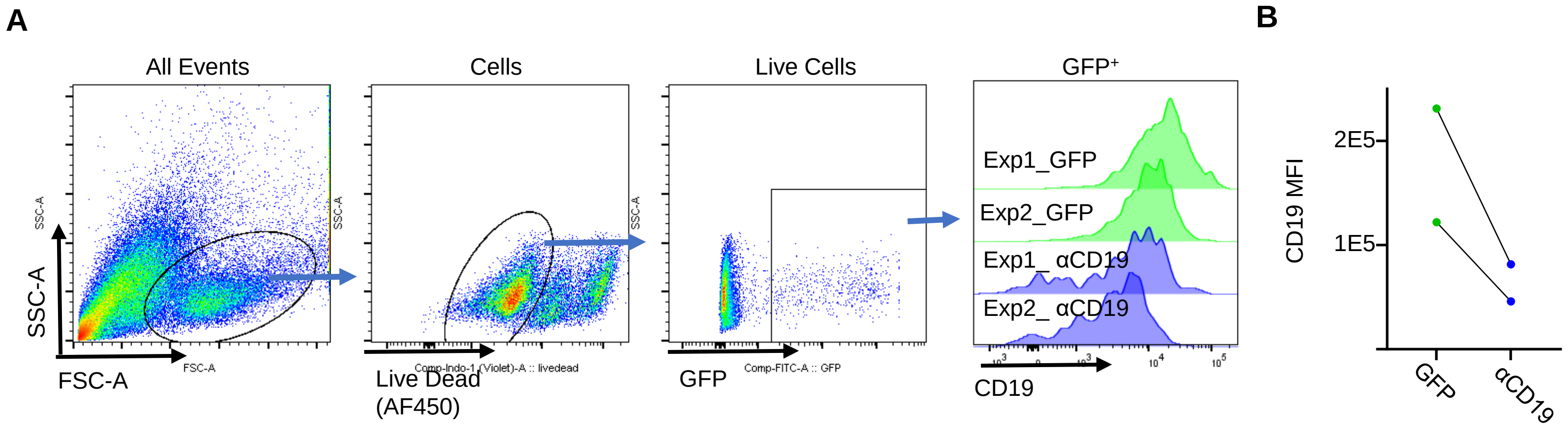


### Supplementary Figure 7

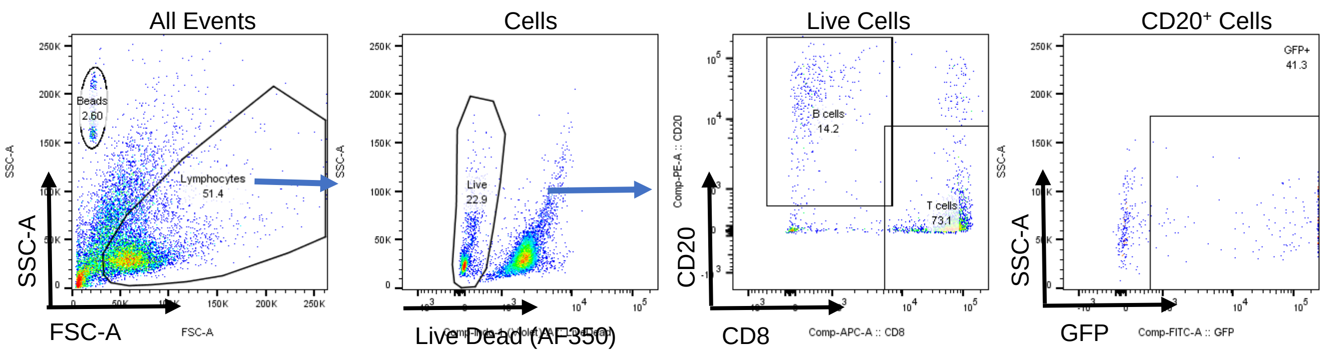


### Supplementary Figure 8

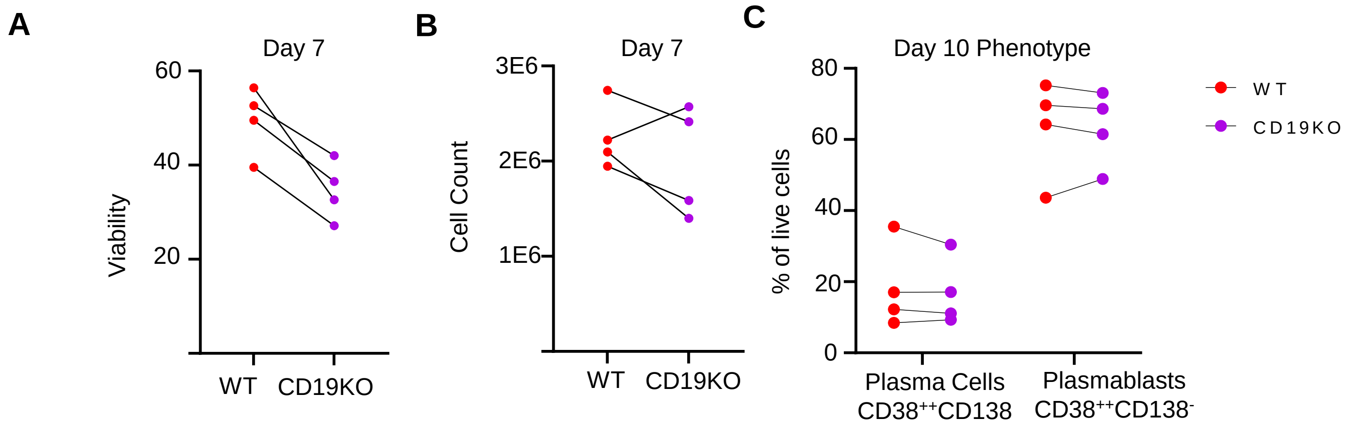

### Supplementary Figure 9

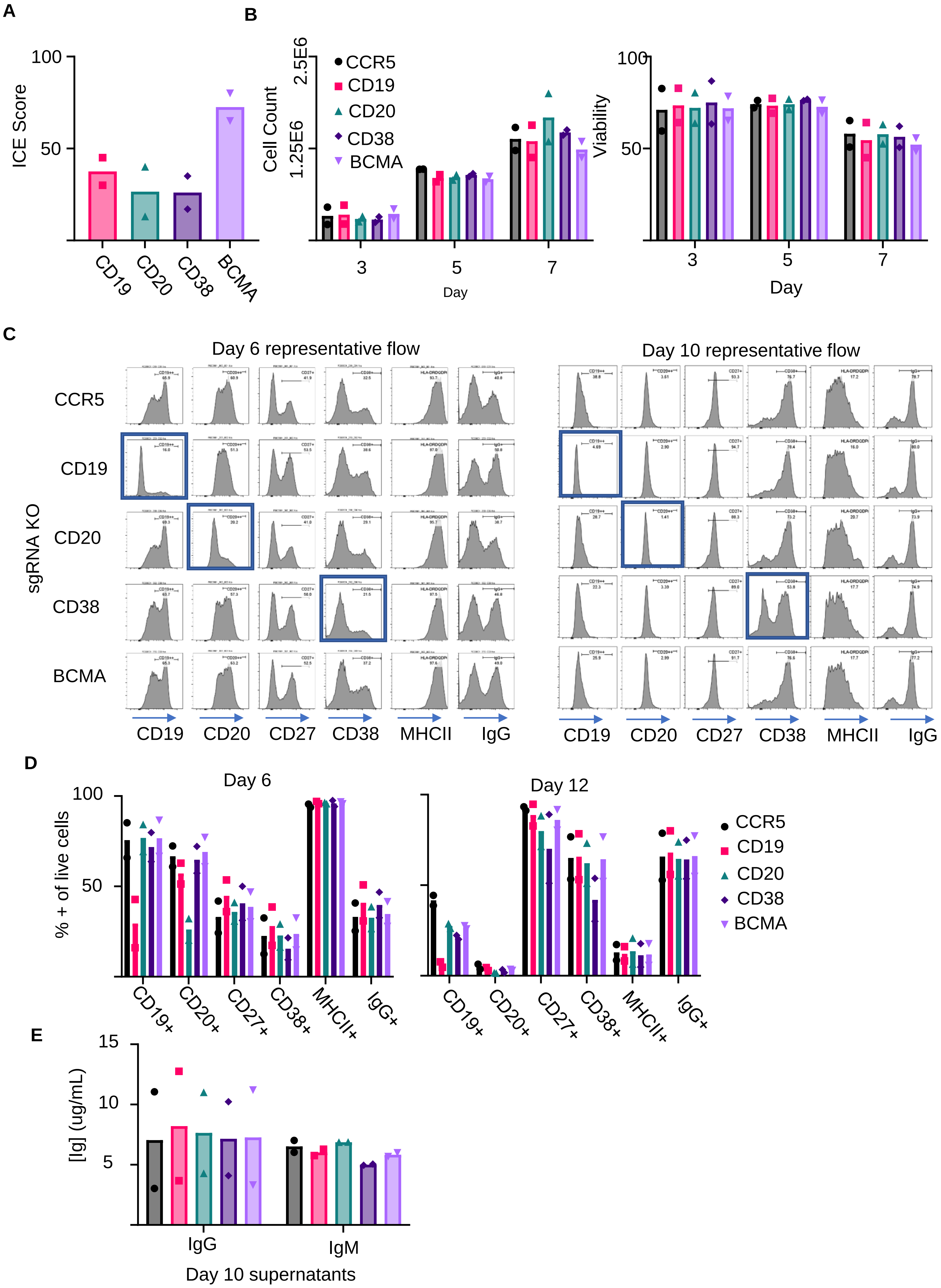

### Supplementals Tables

#### Supplemental Table 1: CRISPR Guide Target Sequences

| Guide Target | Sequence |
|---|---|
| *CCR5* | CAATGTGTCAACTCTTGACA |
| *JCHAIN* | AAGAACCATTTGCTTTTCTG |
| *IgHG1* | TGAGTTTTGTCACAAGATTT |
| Eμ | GTCTCAGGAGCGGTGTCTGT |
| *CD19* | CCTCGGGCCTGACTTCCATG |

#### Supplemental Table 2: Flow Antibody Panels

| **B cell Phenotyping** | | |
|---|---|---|
| **Item** | **Vendor** | **Catalog** |
| PE-Cy7 anti-human CD19 | Biolegend | 302216 |
| PerCP-Cy5.5 anti-human CD38 | BD | BDB551400 |
| PE anti-human CD138 | Biolegend | 356504 |
| Brilliant Violet 605 anti-human CD3 | Biolegend | 317322 |
| GFP | NA | NA |
| BFP | NA | NA |
| Alexa Fluor™ 350 NHS Ester (Succinimidyl Ester) | Fisher | A10168 |
| **K562 Killing Assay** | | |
| **Item** | **Vendor** | **Catalog** |
| BV786 anti-human CD69 | Biolegend | 310932 |
| PE anti-human CD137 | Biolegend | 309804 |
| APC anti-human CD8 | Biolegend | 344722 |
| GFP | NA | NA |
| BFP | NA | NA |
| Alexa Fluor™ 350 NHS Ester (Succinimidyl Ester) | Fisher | A10168 |
| **PBMC Killing Assay** | | |
| **Item** | **Vendor** | **Catalog** |
| BV786 anti-human CD69 | Biolegend | 310932 |
| PE anti-human CD137 | Biolegend | 309804 |
| APC anti-human CD8 | Biolegend | 344722 |
| FiTC anti-human CD14 | Biolegend | 325604 |
| PE-Cy7 anti-human IgM | Biolegend | 314531 |
| PE anti-human CD33 | Biolegend | 303404 |
| Alexa Fluor™ 350 NHS Ester (Succinimidyl Ester) | Fisher | A10168 |
| **Self-Killing Assay** | | |
| **Item** | **Vendor** | **Catalog** |
| BV786 anti-human CD69 | Biolegend | 310932 |
| PE anti-human CD137 | Biolegend | 309804 |
| APC anti-human CD8 | Biolegend | 344722 |
| GFP | NA | NA |
| PE-Cy7 anti-human IgM | Biolegend | 314531 |
| PE anti-human CD20 | BD | 560961 |
| Alexa Fluor™ 350 NHS Ester (Succinimidyl Ester) | Fisher | A10168 |
| **Mouse Tissue Staining** | | |
| **Item** | **Vendor** | **Catalog** |
| APC-Cy7 anti-human CD45 | Biolegend | 368516 |
| Brilliant Violet 605 anti-mouse CD45 | Biolegend | 103155 |
| Alexa Fluor™ 700 anti-human CD4 | Biolegend | 300526 |
| PerCP-Cy5.5 anti-human CD38 | BD | 551400 |
| PE anti-human CD138 | Biolegend | 352306 |
| APC anti-human CD8 | Biolegend | 344722 |
| BV786 anti-human CD3 | Biolegend | 317330 |
| PE-Cy7 anti-human CD19 | Invitrogen | 25-0199-42 |
